# Mapping the Trade-Offs That Shape Mammalian Brain Network Architecture

**DOI:** 10.64898/2026.10.07.757032

**Authors:** Adrian Dendorfer, Alexa Mousley, Francisco Páscoa dos Santos, Duncan E. Astle, Kayson Fakhar

## Abstract

A brain’s wiring must satisfy conflicting pressures, from the outside world (be capable) and from the inside (stay within finite resources). These pressures often have opposing requirements. Prior work studies these trade-offs mostly between 2-3 pressures at a time, and thus cannot say how many pressures shape brain wiring or for which trade-off brains decide. To see all possible trade-offs at once and to uncover the set of most important pressures, we generate 25,000 synthetic brain networks and describe each by 29 network properties. We then place 635 human (ages 6-22) and 224 mammalian (12 taxonomic orders) brain networks in this property space. The space is triangular, and follows three pressures: computational capability, robustness to damage, and metabolic efficiency. No network is optimal at all three, and biological brains cluster in one small niche of this triangle. There, they accept higher metabolic cost to gain both robustness and capability. Development and evolution move brains in the same direction; additionally, human maturation first raises capability, and later optimizes for efficiency. This study underscores the triangular trade-off between metabolic efficiency, computational capacity and robustness, and the narrow region brains occupy in this trade-off triangle, visible only from a multi-property, cross-species perspective.

## INTRODUCTION

The brain is a network, constructed from white-matter tracts that link its regions. This network is called the *connectome*^1^. Its wiring is anything but arbitrary. it is small-world, so any region reaches any other in a few steps^2^. It is modular, so regions fall into communities that talk mostly among themselves^3^. And it has a rich club, a small set of hubs wired densely to one another^4^. Each of these features answers a pressure either from outside the brain - such as the demand to process information and to survive damage - or from inside it - such as a metabolic budget the wiring may not exceed^5–7^.

Wiring that satisfies one of those pressures tends to violate another. Long-range connections are the clearest case: one long edge speeds communication between distant regions, but it is the most expensive wiring a brain can buy, and so buys brain-region’s integration at the price of economy. Modules carry the same tension, because a cleanly divided network supports specialised processing but slows communication between the modules^3,8^. While the precise pressures are not known, candidates are not scarce in prior literature: economy, integration, segregation, robustness to damage, computational capacity and efficient communication have all been proposed^5,6,8–10^. How many of them pull independently, however, and which of them a brain actually resolves, has not been measured. Every claim that a brain is well organised rests on the assumption that we know what it is organised *for*.

A gain towards one pressure is always paid for on another (a *trade-off* ). For instance, because every connection takes space and metabolic energy to build, which are both finite budgets^5^, a brain cannot be perfectly integrated and perfectly metabolically efficient at the same time. Networks that perform the best in a given trade-off construct the *Pareto front*: the set of compromises where nothing can be improved without something else getting worse^11^. The compromise brains take have been analyzed with the Pareto front previously. Avena-Koenigsberger and colleagues introduced the network *morphospace*, a map on which every architecture has a position, and approximated the front that trades information processing against wiring cost^12,13^. Ma and colleagues added segregation as a third objective^14^. Each of these studies, however, fixes its objectives before the analysis starts, so the number of competing demands is set by the authors rather than read off the data. A front drawn through two or three hand-picked measures can therefore show that a trade-off exists, but not how many independent trade-off goals there are, and for which ones the brains decide resolve.

Answering those questions needs many networks that span the entire space of possible trade-offs. These are far more than the few wiring patterns of the connectomes biology provides. Generative network models (GNMs) provide them^15^. A GNM uses a wiring rule that compresses connectome organisation, and then uses that rule alone to build a network edge by edge^16^. At every step the rule scores each potential next edge on two competing terms: the distance the connection would span, which stands for what it costs metabolically, and how many neighbours the two regions already share - a concept observed in real brains termed *homophily*. Two exponents set how heavily each term counts, *η* for the price of distance and *γ* for the reward for shared neighbours. One (*η, γ*) pair is therefore one explicit compromise between cheap wiring and topological affinity, and sweeping the two exponents turns an abstract trade-off into a population of architectures we can measure.

We generated 25,000 networks using a GNM that swept a 50 × 50 grid of (*η, γ*) parameters, and then scored all 25,000 networks on 29 properties. The 29 span every demand that has been proposed for brain wiring, from wiring cost5 and computational capacity^6,17^ to robustness under attack^9^, segregation^3,8^, communication^10^ and critical dynamics. Crucially, they enter the analysis ungrouped, so that whatever structure runs through them is a property of the data and not of a scheme imposed on it. We then measured the same 29 properties on 635 human connectomes aged 6-22 (HCP-D18), on 224 mammalian connectomes from 12 taxonomic orders (MaMI^19^), and on five types of reference networks. Four of them isolate one demand each, and the fifth isolates none. That fifth is an Erdős-Rényi network, which places every connection at random, blind to both distance and topology, and therefore shows one possible architecture that might arise when no demand shapes the wiring. Economy, by contrast, is isolated by a minimum-wiring network, which connects only the 495 closest pairs of regions. It therefore contains no long-range connection and has the smallest total wiring length possible at the given network density. The other three are communication-optimised, each grown to move a signal through the network as well as possible under one assumption about how the signal travels^10^: routing along the shortest path, diffusion as an unguided random walk, or propagation along every path at once.

Principal component analysis of that property matrix defines the *property space*, a morphospace12,20,21 whose axes are trade-offs rather than single properties. A second space, the *GNM morphospace*, keeps *η* and *γ* as its axes, into which all empirical and reference networks are projected. Each connectome therefore carries two coordinates: what it is, and how it was (or could have potentially been) made.

Development and evolution both reshape the connectome, and so offer two natural experiments for watching brains move through this space. During maturation the penalty on long-distance connections weakens, which suggests the economy constraint is progressively relaxed^16,22^. Across mammals, by contrast, global topology stays conserved despite brain volumes that span more than four orders of magnitude^19,23,24^. If competing demands genuinely organise real brains, both kinds of variation should leave structured, low-dimensional traces in the property space, and should be comparable to one another inside it.

Here we therefore ask how many demands genuinely compete in a connectome, and at which compromises mammalian brains settle among them. First, we place all 25,963 networks in the property space and also recover the pairwise and three-way trade-offs that prior work reports. Second, we read the principal components themselves, which resolve into three independent tensions rather than a single front, and so make the space triangular. Third, we group the 29 properties by the tension each one serves, and read every connectome’s position, its donor’s age, and its taxonomic order against that grouping. Finally, we embed every connectome in the GNM morphospace, to ask whether the wiring rule alone carries the same information.

## RESULTS

### Multi-Objective Property Space

Of all the network architectures a brain could in principle have, is the real one special? Answering that needs two things: a space of architectures that are possible, and a way to place real connectomes inside it.

The space of possibilities comes from the GNM sweep (Figure 1B-D; the analysis it feeds is summarised in Figure 1A). Each synthetic network grows edge by edge from a minimum spanning tree to a fixed density of 10 %. At each step, every possible next connection is weighed by two competing terms: the wiring cost and the benefit of the connection. Its score is the distance it would span, raised to the power *η*, times the homophily of the two regions it would join, raised to the power *γ* (see Section 4.2). Sweeping *η* ∈ [−8, 3] and *γ* ∈ [−0.1, 1] on a 50 × 50 grid with ten replicates per cell gave 25,000 networks. The two exponents visibly change what gets built (Figure 1D): with *η <* 0 edges stay short and locally clustered and wiring cost is low (locations A and E), and with *η >* 0 long edges spread across the whole brain and wiring cost is high (locations B and D). Thus, the sweep supplies a systematic population of architectures, each built from a different trade-off, against which real connectomes (or further synthetic networks) can be measured.

**Figure 1:**
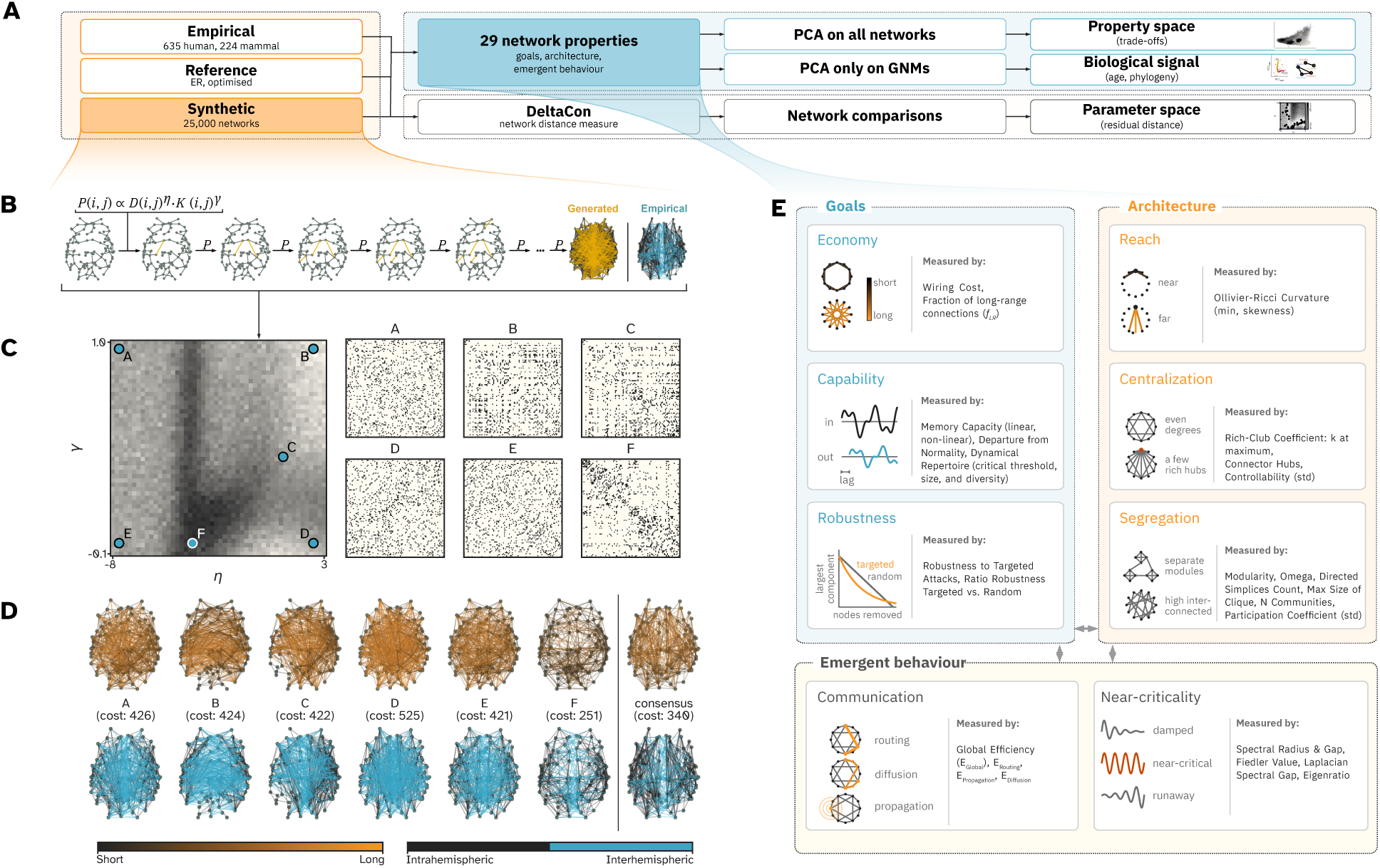
Analysis pipeline, the generation of the synthetic networks with the GNM parameter sweep, and an overview over the 29 measured properties. (A) Analysis pipeline, where three sets of networks enter: empirical connectomes (635 human, 224 mammal), reference networks (Erdős-Rényi, minimum-wiring network, communication-optimised), and synthetic networks (25,000 GNMs). Along the upper path, every network is characterised by the same 29 properties, grouped into goals, architecture, and emergent behaviour. A PCA on all networks then gives the property space, and a PCA only on the GNMs, into which the connectomes are projected, serves for the age and phylogeny analyses of the biological signal. Along the lower path, the network distance measure DeltaCon compares each connectome with the synthetic networks, placing it in the (*η*, *γ*) parameter space with a best-fit pair and a residual distance. Panels B-D expand the synthetic networks, panel E the 29 properties. (B) Creation of the synthetic networks. This is an iterative process: starting from a minimum spanning tree seed (grey edges), edges (orange) are added one at a time with probability *P* (*i, j*) ∝*D*(*i, j*)*^η^ K*(*i, j*)*^γ^*, where *D* is the distance and *K* the homophily between regions *i* and *j*, until the empirical density is reached. Right: one generated network (orange) next to an empirical consensus connectome (blue). (C) The left side shows the (*η*, *γ*) parameter space (*η*∈[ −8, 3], *γ*∈ [ − 0.1, 1]) with six illustrative parameter combinations (marked A-F). The background colour denotes similarity to the HCP-D consensus connectome (dark: more similar). The right plots show the binary adjacency matrices of (A-F); dark entries depict a present connection. (D) The same six networks (A-F) and the human (HCP-D) consensus connectome rendered on Schaefer-100 coordinates, with wiring cost below each. In the top row, edge colour encodes connection length (dark: short; orange: long); in the bottom row it encodes hemisphere (dark: intrahemispheric; blue: interhemispheric). (E) The 29 properties, grouped by their different associations: goals, architecture, and emergent behaviour.

Into this population we projected four other sets of networks: empirical connectomes from humans (*n* = 635, HCP-D^18^; Section 4.1) and from other mammals (*n* = 224, MaMI ^19,24^; Section 4.1); one structural baseline, namely 100 Erdős-Rényi random networks; and networks that were optimized for one specific purpose. This includes a minimum-wiring network that connects only the 495 closest pairs of regions (Section 4.2) and three communication-optimised reference networks, one each for routing, diffusion, and propagation (Section 4.2). Each of these three was grown to be as good as possible at one single way of moving a signal10. Routing sends the signal along the shortest path. Diffusion lets it wander as a random walk. Propagation broadcasts it along all paths at once. They show what an architecture looks like when one communication regime is optimised without compromise, and so bracket the biological networks from a direction the GNM sweep does not cover.

Every network was then characterised by the same 29 structural, dynamic, and computational properties (Figure 1E; see Sections 4.3 and 4.3). Together, these 29 properties cover the demands that have been proposed for brain wiring. We analysed the property space first without any grouping of the properties, so that the structure among them would emerge from the data rather than from a scheme imposed on it. The grouping drawn in Figure 1E, grouping the properties into goals, architecture, and emergent behaviour, therefore does not enter the PCA. Instead, each property was assigned by hand to one of the groups (Section 4.3). For this reason, we introduce the grouping only once the geometry of the property space has been described. Principal component analysis (PCA) of the resulting property matrix defines a low-dimensional property space (see Section 4.4 and Figure 2). PC1 accounts for 57.49 % of total variance, PC2 for 14.65 %, and PC3 for 4.51 %.

**Figure 2:**
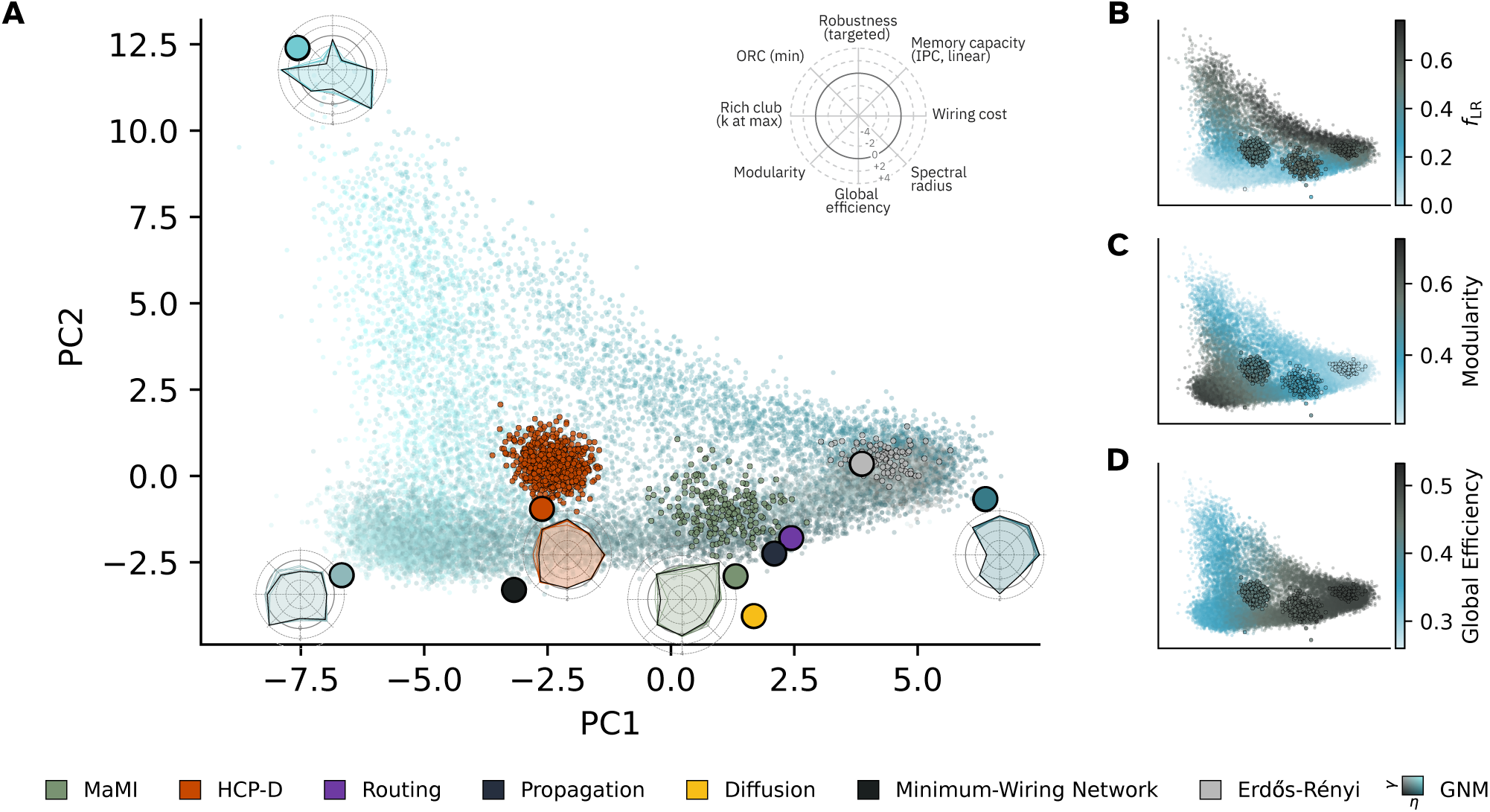
Property space of all 25,963 networks, with per-location profiles. (A) PC1-PC2 scatter of all 25,000 GNM networks (teal cloud; colour encodes *η* and *γ* values, going from low *η* and high *γ* values in black, via high *η* and low *γ* values in dark blue, to high *η* and *γ* values in light blue; see legend), HCP-D connectomes (orange, *n* = 635), MaMI connectomes (olive, *n* = 224), structural baselines (Erdős-Rényi random networks, grey; minimum-wiring network, black), and three communication-optimised reference networks, each grown to be as good as possible at one signalling model alone (routing along the shortest path: purple; propagation along all paths at once: dark navy; diffusion by random walk: yellow). PC1 accounts for 57.49 % and PC2 for 14.65 % of total variance. Radar insets show exemplary property profiles: robustness to targeted attacks, linear memory capacity (IPC_1_), wiring cost, spectral radius, global efficiency, modularity, the rich-club coefficient (*k* at maximum), and minimum ORC. Values are standardised. The five radar plots are drawn for five networks: the three most extreme synthetic (GNM) networks, and one connectome from each of HCP-D and MaMI. (B-D) The same scatter coloured by the fraction of long-range connections *f*_LR_ (B), modularity (C), and global efficiency (D).

The radar insets give an intuition for how the space varies from place to place (Figure 2A). Each plots the same eight properties, one standing in for each group, at one of five landmark locations: three at the most extreme GNM networks, which mark the corners the cloud spans, and one connectome from each of the two empirical datasets. Six of the eight properties are explained in Supplementary Note 2; all 29 are defined in Supplementary Note S1.

Both HCP-D and MaMI connectomes cluster in a bounded region of the property space, entirely enclosed by the GNM cloud (Figure 2A). The propagation- and routing-optimised networks fall very close to the MaMI cluster. The diffusion-optimised network falls outside the GNM cloud altogether and marks, together with the minimum-wiring network, the low-PC2 extreme. The Erdős-Rényi networks mark a further extreme, occupying the regime of minimum PC1 values, and the GNM networks in that region are mostly generated by low *η* and low *γ*. Empirical connectomes therefore converge on one specific region of this multi-objective property space, rather than scattering across it or occupying an extreme pole.

Sitting in the midst of the cloud, empirical connectomes do not maximise any single property. They hold moderate values on several objectives at once. This indicates that the empirial connectomes are trading-off different competing demands. Three such properties are shown in Figure 2B-D and further ones in Supplementary Fig. S4. Strikingly, this convergence holds across mammals spanning more than four orders of magnitude in brain volume23: a small bat and a large ungulate land in the same region. Brain-like wiring constraints thus appear to drive connectomes toward a shared multi-objective optimum, largely independent of phylogenetic history. Within that region, however, the two empirical datasets form adjacent rather than intermingled clouds, and we do not read the offset as a biological boundary. HCP-D and MaMI differ in acquisition (in vivo against ex vivo), in parcellation (Schaefer-100 against k-means clustering of fibre endpoints), and in tractography pipeline, and any of these could displace one cloud relative to the other.

### Recovering Established Trade-Offs: Pareto-Optimal Frontiers

If biological connectomes converge on one bounded region, what compromise does that region represent? Prior work reports a range of pairwise trade-offs between network properties^5,13,14^. Before reading the space as a whole, we check that two of those known trade-offs reappear across the 25,000-network population, and locate the empirical connectomes on each.

Global efficiency and modularity define a strong negative Pareto front (Figure 3A). The Pareto front separates unachieveable combinations from combinations that are reachable by networks. This trade-off is the classic tension between integration and segregation^8^: adding long-range connections raises global efficiency at the expense of community structure. In the GNM cloud, that means moving toward the greyer colours. Empirical connectomes sit at a compromise position (medoid of HCP-D: modularity ≈ 0.50, global efficiency ≈ 0.43; medoid of MaMI: ≈ 0.40 and 0.47). The routing-optimised reference network, by contrast, anchors the high-efficiency, low-modularity extreme (modularity ≈ 0.22, global efficiency ≈ 0.53).

**Figure 3:**
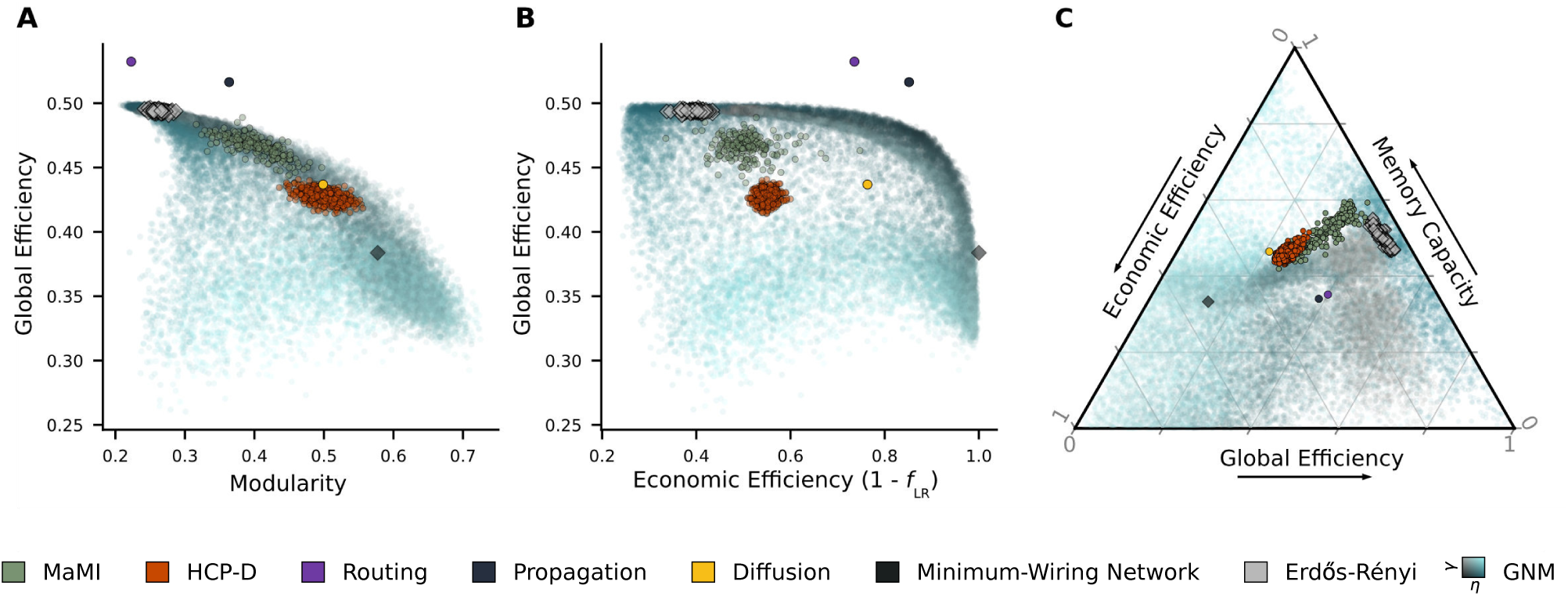
Pairwise and three-way trade-off plots across all networks. All 25,000 GNM networks (teal), empirical connectomes (orange: HCP-D; olive: MaMI), communication-optimised reference networks (routing: purple; propagation: dark navy; diffusion: yellow), a minimum-wiring network (dark diamond), and structural baselines (Erdős-Rényi: grey diamonds) are shown in every panel. (A) Global efficiency vs. modularity. A clear negative Pareto front separates achievable from unachievable combinations; empirical connectomes cluster at a compromise of high-to-moderate global efficiency and low-to-moderate modularity, while routing-optimised networks anchor the high-efficiency, low-modularity extreme. (B) Economic efficiency (1 − *f*_LR_) vs. global efficiency. A second negative front; empirical connectomes cluster at an interior compromise, while the routing- and propagation-optimised networks reach the highest global efficiency and high economic efficiency, and the ring lattice anchors the maximally economical, lower-efficiency corner. (C) Ternary Pareto triangle with vertices at economic efficiency (1 − *f*_LR_, left), global efficiency (right), and memory capacity (top). Each network is positioned by its three rank-normalised property values, closed to sum to one, so that all vertices are reachable (see Section 4.5). The memory-capacity axis uses the standalone memory capacity (i.e. forgetting-curve) estimate rather than the IPC_1_ used elsewhere.

Economic efficiency and global efficiency define a second negative front (Figure 3B). Economic efficiency defined as 1 − *f*_LR_, where *f*_LR_ is the fraction of long-range edges, and thus 1 − *f*_LR_ represents the degree to which a network minimizes wiring cost (it is 1 when no connection is long-range, and 0 when all connections are long-range). The trade-off is the foundational tension between metabolic cost and integration^5,13^; conserving wiring by forgoing long-range connections (i.e. raising economic efficiency) lowers global efficiency. Both empirical connectomes and the GNM cloud lie at an interior compromise, combining low-to-moderate economic efficiency with high-to-moderate global efficiency (HCP-D: 1 − *f*_LR_ ≈ 0.55, global efficiency ≈ 0.43; MaMI: ≈ 0.51 and 0.47). The routing-and propagation-optimised networks reach higher global efficiency (≈ 0.53 and 0.52), above the cloud, while the minimum-wiring network anchors the maximally economical, low-efficiency corner (1 − *f*_LR_ = 1.00, global efficiency ≈ 0.38).

Third, the constraints can be summarised into a three-way Pareto triangle with vertices at economic efficiency (1 − *f*_LR_, which closely proxies absolute wiring cost; Fig. 7A), global efficiency, and memory capacity (Figure 3C)^11^. Each axis is rank-normalised so that all vertices are reachable (see Section 4.5). The memory-capacity axis uses the standalone forgetting-curve estimate, i.e. the summed linear-capacity-versus-delay curve whose per-delay profile is plotted in Fig. 7C, not the IPC_1_ that defines the property space. The two rank networks alike, and only the relative ordering matters here (see Section 4.3). The GNM networks fill the simplex unevenly. They are densest toward the economy and global-efficiency vertices, and sparse toward the memory-capacity vertex, which no network approaches. Both HCP-D and MaMI cluster near the edge spanning high global efficiency and high memory capacity, far from the economy vertex. This points to stronger pressure toward computation and integration than toward metabolic economy. It also confirms convergence on the same compromise across mammalian evolution and human development. The ternary floor shows only the *relative* balance among the three objectives and discards how high a network scores on them in total. The triangular-prism rendering of Supplementary Fig. S5 restores that magnitude as an orthogonal height axis, each point’s height being its summed rank across the three axes (a rotating version is available at https://github.com/ad045/14_4D_property_space/blob/main/additional_material/trade-off_3D.gif). There, GNM networks with exceptionally low *γ* reach the highest total scores, higher than any empirical connectome. The empirical connectomes themselves sit at intermediate heights, despite lying close to the memory-capacity frontier. Among the reference networks, the propagation-optimised network is both the most total-optimal and the most central on the floor (i.e. the most evenly balanced across the three axes). That empirical connectomes are not the most total-optimal on these three objectives suggests they also trade off against objectives outside this triangle.

### Competing Objectives and the Property Scheme

As shown in the previous section, two fronts and the simplex recover trade-offs that prior work reports. However, both prior work and Section 2.2 fix their objectives in advance. Thus, they cannot say how many objectives primarily shape the property space, and which ones they are. In the following, the principal components were used to choose the objectives (Figure 4A).

**Figure 4:**
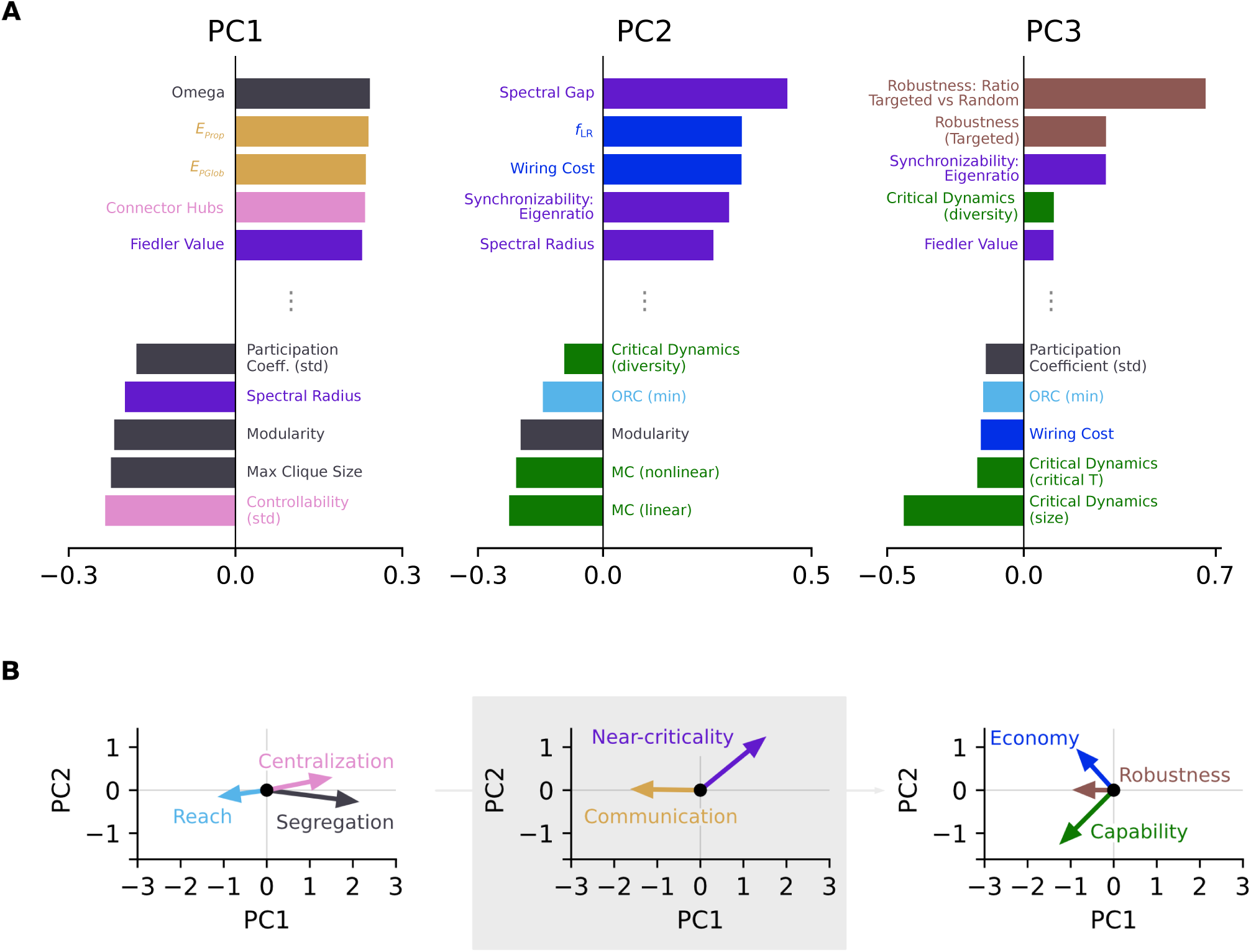
Property loadings on the first three principal components, and the group directions they define. (A) The five largest positive and negative property loadings on PC1, PC2, and PC3. Bars are coloured by group (the three goals (economy, capability, robustness), the three architectural settings (reach, segregation, centralization), and the two emergent behaviours (communication, near-criticality)). Loadings are shown for each property in its native orientation, that is before the within-group sign alignment described in Section 4.4, so that every bar reflects the property as conventionally defined. The omega bar, for example, reflects small-worldness itself, not its complement. Sign-inverting a property before the PCA only flips the sign of its own loading, and leaves the axes, the explained variance, and every network’s position unchanged, so this is purely a matter of presentation. Because the sign of each principal component is itself arbitrary, the orientation of the loading bars is independent of the arrow orientation in (B); only the groupings and oppositions, not the absolute left-right direction, should be compared across the two panels. (B) Group directions projected as vectors into the PC1-PC2 plane, separated by level: *left*, the three architectural settings (reach, segregation, centralization); *centre*, the two emergent behaviours (communication, near-criticality); *right*, the three goals (economy, robustness, capability). Each arrow is the summed biplot loading of that group’s properties. Because a group’s direction is the mean of its members’ loadings, this panel uses the sign-aligned (inverted) properties, so that all members of a group point the same way and their mean direction is meaningful rather than a near-cancellation of anti-correlated members.

PC1 separates small-worldness (omega), global efficiency, and propagation efficiency from modularity, maximum clique size, and controllability (std). PC2 separates the spectral gap, the fraction of long-range connections, and wiring cost from memory capacity (linear and non-linear) and modularity. PC3 separates the targeted-to-random robustness ratio, robustness against targeted attacks, and synchronizability from critical dynamics (critical size and threshold) and wiring cost. In broad brush strokes, PC1 reflects the tension between integration and segregation, PC2 reflects the trade-off between wiring cost and computational capacity, and PC3 sets robustness against all other objectives.

Of the three PCs, only PC2 and PC3 carry trade-offs between goals. PC1 instead records the balance between integration and segregation, which are architectural styles that serve goals rather than goals in themselves. The goals that remain are economy, capability, and robustness. Because these three pull against one another along two independent components, they span a triangle in the 2D projection of the property space.

After identifying the three main goals, we investigated how the 29 properties relate to them. We manually defined the following structure (Figure 1E):

The three *goals* (economy, capability, robustness) are realized by three *architectural* settings (reach, segregation, centralization). From the architecture, two *emergent behaviours* (communication, near-criticality) arise that also contribute to the goals. To compare this manual grouping to the data, we projected each group’s summed loading direction into the PC1-PC2 plane, treating the three levels separately (Figure 4B).

The summed loading directions of the *goals* (Figure 4B, right) reproduce the trade-off on PC2: economy points toward positive PC2, and capability toward negative PC2. The robustness arrow carries almost no PC2 component and is much shorter than the other two, so most of its weight lies in higher components, consistent with its loading on PC3. The plots for the *architecture* (Figure 4B, left) and the *emergent behaviours* (Figure 4B, middle) provide further information for interpreting the property space.

### Developmental Maturation and Phylogenetic Information

Is a connectome’s position within this landscape biologically meaningful, or merely incidental? To find out, we tested whether position tracks two independent biological variables: human development and mammalian phylogeny.

Such a test needs axes that the biological data had no hand in defining. We therefore fitted a second PCA on the GNM networks alone and projected the connectomes into it afterwards, naming its axes PC1_GNM_ and PC2_GNM_ to distinguish them from the joint property space of Section 2.1. Comparing the two confirms that the few hundred empirical, optimised, and baseline networks barely move axes dominated by 25,000 GNM networks. Both GNM-fitted axes carry biologically interpretable signals (Figure 5, see Section 4.6).

**Figure 5:**
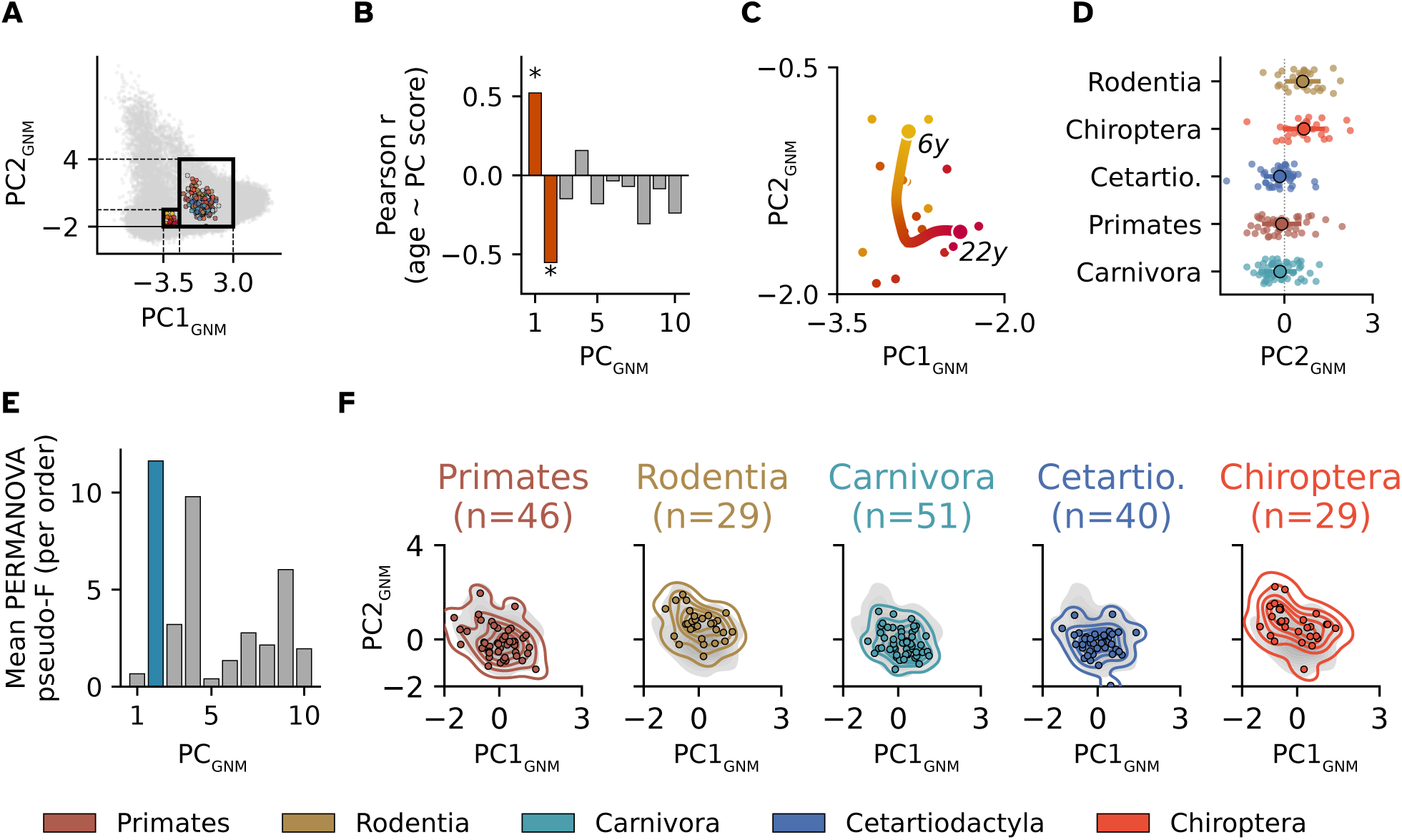
Developmental and phylogenetic structure in the GNM-fitted property space. All panels use the property space defined by PCA fit on GNM networks. (A) PCA fit only on GNM networks, including MaMI connectomes and the HCP-D consensus networks. Black rectangles mark the regions that are magnified in the following panels. (B) Pearson *r* between age and PC_GNM_ score for components 1-10, computed across the 17 per-year HCP-D consensus observations. PC1_GNM_ (*r* = 0.521, *p* = 0.032) and PC2_GNM_ (*r* = −0.552, *p* = 0.022) are the only significant age correlates. (C) Smoothed developmental trajectory (cubic smoothing spline, *λ* = 10) of the 17 consensus connectomes, coloured from age 6 (yellow) to 22 years (red), with the point of maximum curvature at 16.0 years. (D) Mean PERMANOVA pseudo-*F* statistic per component, averaged across the five mammalian orders with *n >* 20. PC2_GNM_ yields the highest mean phylogenetic separation (*F̂* = 11.63), followed by PC4_GNM_ (*F̂* = 9.79). (E) Per-order PC1_GNM_-PC2_GNM_ kernel density plots (Primates, Rodentia, Carnivora, Cetartiodactyla, Chiroptera), with the combined density of the selected orders in grey for reference. (F) The PC2_GNM_ scores by order. (G) Pairwise Mahalanobis distances between the five taxonomic orders with *n >* 20 in PC1_GNM_-PC2_GNM_ space. (H) Graph of non-significant pairwise PERMANOVA comparisons (*p >* 0.05): edges connect orders that are not statistically separated, revealing two clusters, (Primates, Carnivora, Cetartiodactyla) and (Rodentia, Chiroptera). (I) Phylogenetic divergence times (in million years) for the same order pairs, retrieved from TimeTree26; pairs with divergence times of 94 MYA (million years ago) are excluded for clarity.

A non-linear developmental trajectory with a late-adolescent inflection

We first built one consensus network from all HCP-D individuals sharing the same integer age, yielding 17 per-year consensus observations spanning ages 6-22 years (Section 4.1). Across these observations, age correlates significantly with both PC1_GNM_ (*r* = 0.521, *p* = 0.032) and PC2_GNM_ (*r* = −0.552, *p* = 0.022), and with no other component (all |*r*| ≤ 0.31; Figure 5B). The developmental signal is therefore concentrated in the first two components of the GNM-fitted property space.

The smoothed trajectory through PC1_GNM_-PC2_GNM_ space is markedly non-linear (Figure 5C; Section 4.6). Between ages 6 and roughly 16 years, displacement runs mostly along PC2_GNM_; the direction of travel then rotates, and the trajectory runs primarily along PC1_GNM_. Curvature analysis of the spline path places this inflection at 16.0 years, which corresponds to the transition from mid- to late adolescence, and is stable across all smoothing parameters *λ* ≥ 5 (Supplementary Note S8). In this PCA, lower PC2_GNM_ values correspond to higher computational capacity and higher PC1_GNM_ values to stronger integration (Supplementary Fig. S6), so the connectome first shifts toward computational capacity and then reorients toward integration.

Phylogenetic separation is concentrated in PC2_GNM_

Across MaMI mammals, we ran a permutational multivariate analysis of variance (PERMANOVA)^25^ on each component, averaging the pseudo-*F* statistic across the five orders with *n >* 20 (Carnivora: *n* = 51; Primates: *n* = 46; Cetartiodactyla: *n* = 40; Rodentia: *n* = 29; Chiroptera: *n* = 29). PC2_GNM_ scores highest (*F̂* = 11.63), followed by PC4_GNM_ (*F̂* = 9.79). All remaining components separate the orders far less well (*F̂* ≤ 3.20; Figure 5D). Per-order density plots show that the orders occupy different parts of the space, although their clouds do overlap (Figure 5E). What separates them is the location of their density peaks along PC2_GNM_, not a clean partition of the plane: Rodentia peak at the highest PC2_GNM_ values and Primates at the lowest (Figure 5F), consistent with compact, locally embedded wiring in small-brained mammals^19,23^. Notably, PC2_GNM_ carries both the strongest developmental and the strongest phylogenetic signal. The same dimension of network organisation therefore varies systematically across developmental time and across evolutionary history.

To quantify the separation between mammalian orders we compared pairwise PERMANOVA (999 permutations) with Mahalanobis distance (Section 4.6; Table 1). Together they reveal a two-cluster structure among the five main orders (Figure 5G-H). Primates, Carnivora, and Cetartiodactyla form one cluster, with small Mahalanobis distances (*d_M_* = 0.22) and non-significant PERMANOVA comparisons (*p >* 0.05). Rodentia and Chiroptera form a second (*d_M_* = 0.18), likewise not separated internally, but clearly separated from the first (all *p* = 0.001, *F >* 10). The split is driven predominantly by PC2_GNM_, identified above also to be the strongest axis of developmental signal. The separation does not recapitulate the phylogenetic tree, but follows more the level of cortex folding or the brain volume.

**Table 1:** Pairwise PERMANOVA (999 permutations) in the GNM-fitted property space (PC1_GNM_–PC2_GNM_). *F* : pseudo-*F* statistic. *p*: permutation *p*-value. Cohen’s *d*: Mahalanobis distance between group centroids using pooled covariance (multivariate effect size). Significance: *** *p <* 0.001, ** *p <* 0.01, * *p <* 0.05, ns *p* ≥ 0.05.

| **Order A** | **Order B** | *F* | *p* | Cohen's *d* |
| --- | --- | --- | --- | --- |
| Primates | Rodentia | 10.68 | **0.001 \*\*** | 0.03 |
| Primates | Carnivora | 0.65 | 0.534 ns | 0.22 |
| Primates | Cetartiodactyla | 0.12 | 0.889 ns | 0.00 |
| Primates | Chiroptera | 10.57 | **0.002 \*\*** | 0.15 |
| Rodentia | Carnivora | 17.09 | **0.001 \*\*** | 0.19 |
| Rodentia | Cetartiodactyla | 15.17 | **0.001 \*\*** | 0.03 |
| Rodentia | Chiroptera | 0.27 | 0.728 ns | 0.18 |
| Carnivora | Cetartiodactyla | 0.61 | 0.565 ns | 0.24 |
| Carnivora | Chiroptera | 17.05 | **0.001 \*\*** | 0.37 |
| Cetartiodactyla | Chiroptera | 14.19 | **0.001 \*\*** | 0.16 |

### GNM Morphospace

So far the GNM sweep has served as a backdrop. We now ask a sharper question of the rule itself: can cost plus homophily actually build a network where real brains sit? Answering it means reading the second morphospace, the one whose axes are *η* and *γ* rather than network properties. We located each empirical and reference connectome in it by assigning it the parameters (*η*^*, γ*^) of the single most structurally similar GNM network, measured by DeltaCon27 (Figure 6, heatmaps; see Section 4.7). To summarise a whole dataset we took its medoid, the best-fit coordinate that corresponds to an actual network in that dataset.

**Figure 6:**
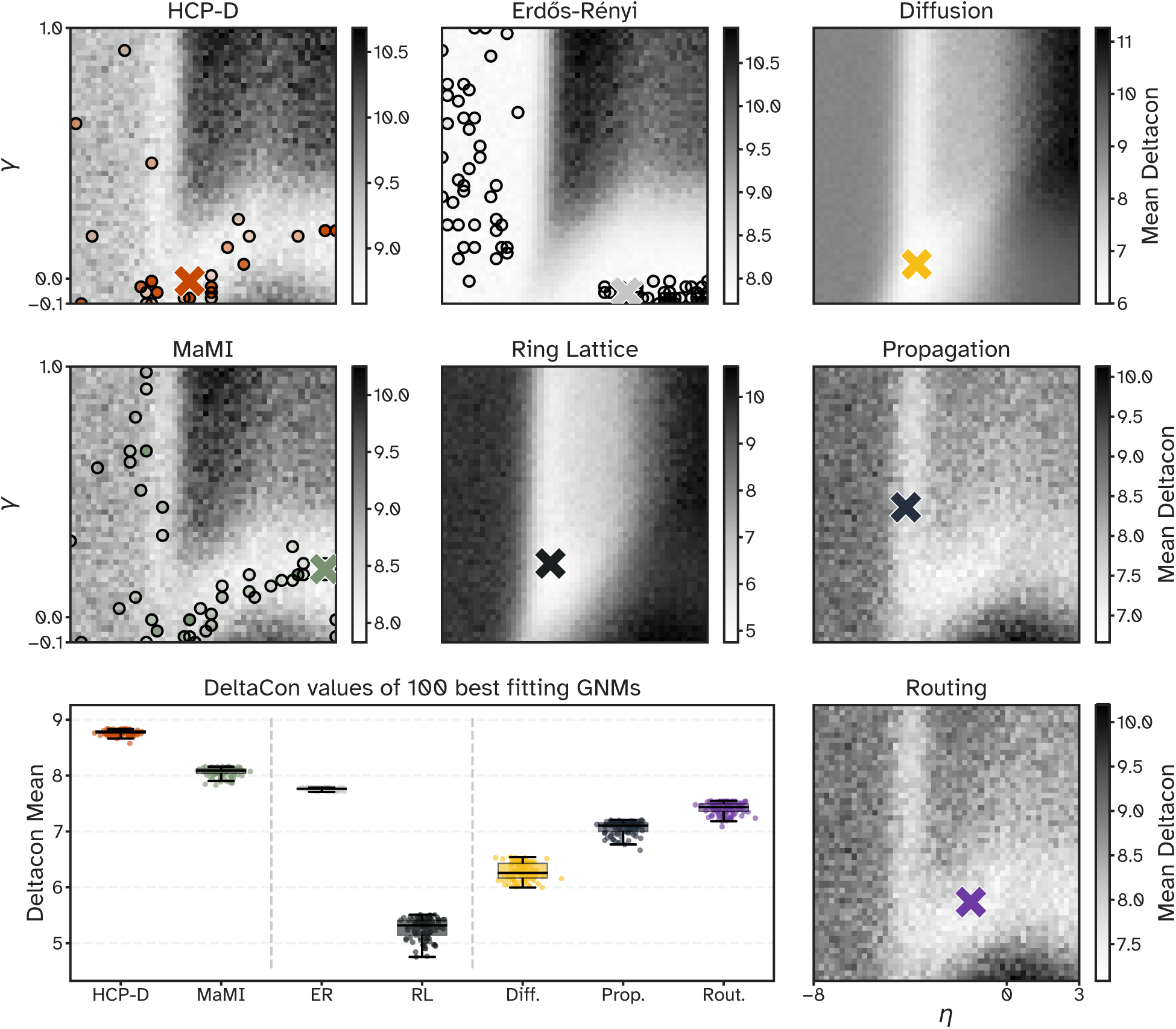
Empirical and reference connectomes embedded in the GNM morphospace via DeltaCon. Each heatmap panel shows the mean DeltaCon distance *d̂D*(*η, γ*) from every (*η, γ*) grid point to the target dataset, averaged across stochastic realisations and across the dataset’s connectomes (lighter = lower = better structural match). Dots mark the best-fit coordinate of each individual connectome, i.e. the (*η, γ*) of the single synthetic network most similar to it; for HCP-D, MaMI, and ER, overlapping semi-transparent dots make recurrently selected coordinates appear darker, indicating local density. Note that, because the dots derive from single closest networks while the surface is realisation-averaged, a dot need not lie on the lightest cell. The large cross in each panel marks that dataset’s medoid, i.e. the best-fit coordinate minimising the total distance to all other best-fit coordinates in the dataset (Section 4.7); in the panels holding only one network (minimum-wiring network, diffusion, propagation, routing) it sits on that network’s single dot. The boxplot (bottom left) shows the *residual distance* per dataset: the distribution of *d̂D* over the 100 lowest-distance grid cells. The biological connectomes (HCP-D, MaMI) carry the highest residual distances.

**Figure 7:**
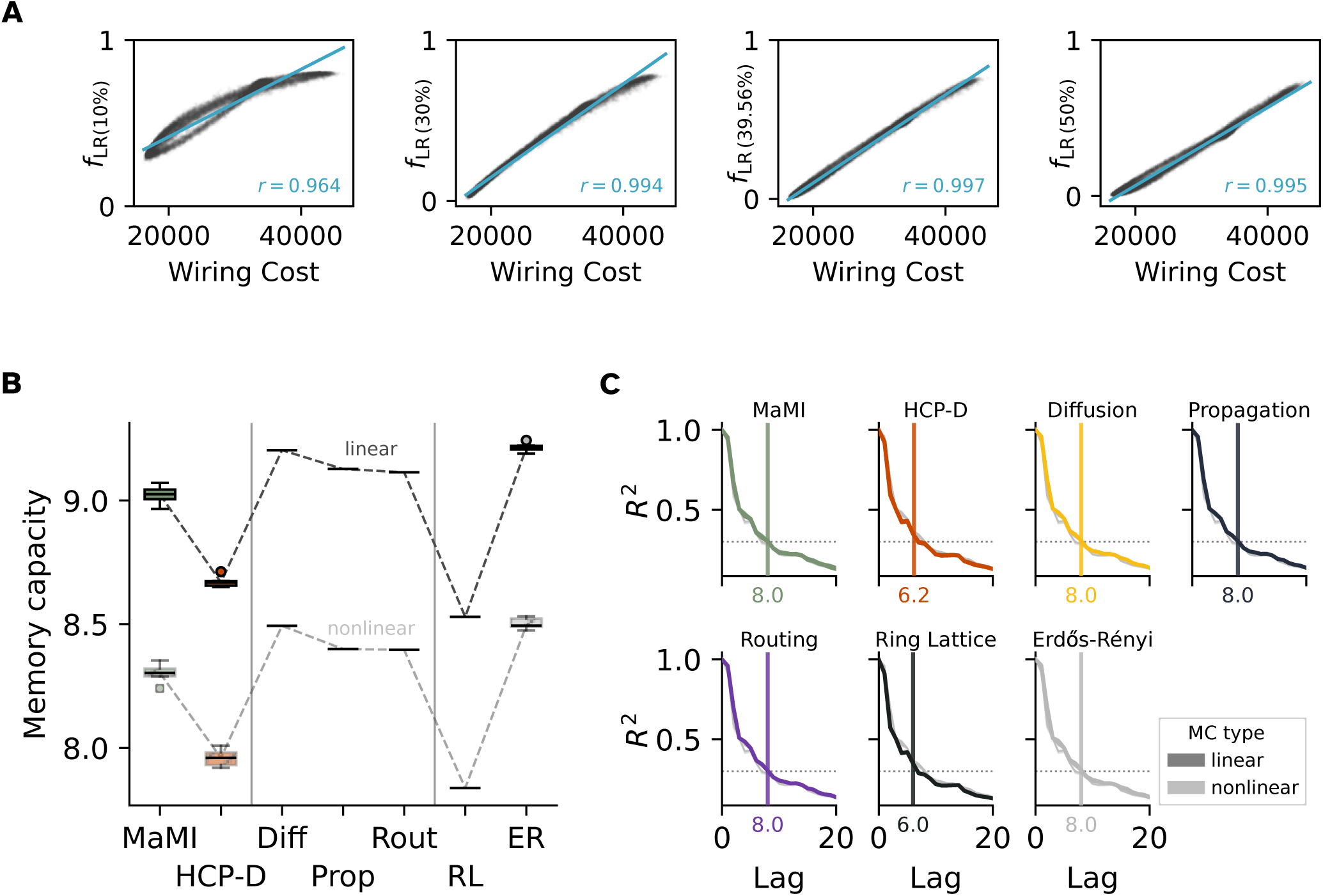
The experiments behind the economy and memory readouts. (A) Fraction of long-range connections *f*_LR_ against absolute wiring cost *W* = Σ*ij a_ij_d_ij_*for the 25,000 GNM networks, at four candidate percentile thresholds of the empirical edge-length distribution (10th, 30th, 39.56th, 50th; left to right). Blue line, linear fit; *r*, Pearson correlation. (B) Mean linear (dark) and non-linear (light) memory capacity per dataset: MaMI, HCP-D, the diffusion-, propagation- and routing-optimised networks (Diff, Prop, Rout), the minimum-wiring network (MW), and Erdős-Rényi (ER). Boxes give the spread across trials; dashed lines join the dataset means. (C) Linear (dark) and non-linear (light) capacity against input delay for lags 0-20, one panel per dataset. The dotted line marks the fixed *R*^2^ = 0.3 criterion, and the vertical line and number give the lag at which each dataset falls below it.

Connectomes from the HCP-D dataset cluster tightly, best represented by their medoid at *η* ≈ −3.06 and *γ* ≈ −0.01; the unexpected negative *γ* may reflect preprocessing choices. MaMI best-fit coordinates span a broader region, either a vertical band around *η* ≈ −5 or a diagonal strip from (*η, γ*) ≈ (−4, −0.1) to (3, 0.2), which likely reflects the greater morphological diversity across 224 species or artefacts of the less established preprocessing. Its medoid at (*η, γ*) ≈ (2.55, 0.19) carries a positive *η*, which would mean connection probability *rising* with distance. That is not a quirk of the medoid: 62.5 % of MaMI connectomes have a best fit with *η >* 0, against 6.1 % of HCP-D connectomes. A majority of the mammalian dataset therefore fits best in a regime no brain can occupy, which we read as a signature of the MaMI preprocessing rather than of mammalian wiring. The three communication-optimised networks land in distinct regions, consistent with what each was grown to do. The routing-optimised network, which favours shortest paths, sits at *η* ≈ −1.49 and moderate *γ* ≈ 0.21. That is the weakest distance penalty of the three, as shortest paths need long edges. The diffusion- and propagation-optimised networks sit further left, at *η* ≈ −3.73 and −4.18. Of those two, propagation carries by far the higher *γ* (0.44 against 0.06), reflecting the stronger homophily that dense multi-scale interconnection requires.

A best fit, however, need not be a good fit. Even at the best-fitting parameters the *residual distance* remains substantial (Figure 6, boxplot; the realisation-averaged distance over the 100 best-fitting parameter combinations). It is the biological connectomes - not the random or regular baselines - that retain the highest residual distances (HCP-D: 8.77 ± 0.05; MaMI: 8.07 ± 0.08). They exceed the minimum-wiring network (5.26 ± 0.19), the diffusion-optimised network (6.28 ± 0.16), and the propagation-optimised network (7.06 ± 0.12). Some of that gap is unsurprising, because an empirical connectome carries measurement noise and inter-individual variability that a rule-generated network does not. But the minimum-wiring network and the three optimised networks are not GNM products either, and the rule still reproduces them more closely than it reproduces a brain. The cost-homophily rule therefore captures the broad topological character of biological connectomes, but not their specifics: plausible hub placement, the spatial distribution of long-range connections, or fine-grained community structure^28^.

The same two-dimensional (*η, γ*) space also compresses away the variation that distinguishes mammalian orders. PERMANOVA on the best-fit coordinates yields non-significant pairwise separations between most orders (Supplementary Note S9), in contrast to the clear separation seen in the property space.

Most of the GNM morphospace is in any case empty, and the regions the empirical connectomes occupy are sparse, as reported before for morphospaces generally12,20.

## DISCUSSION

Real brains occupy a small, non-random region of the architectures a brain-like wiring rule can build. The axes of that region read as competing demands rather than as independent properties. Earlier morphospaces reached a similar conclusion from only two or three graph metrics^5,12,13^. This study, spanning 29 properties, resolves the compromise into three distinct axes instead: a dominant integration-segregation axis, a second pitting wiring cost against computational capacity, and a third trading robustness against every other domain. Because biological wiring sits in the interior rather than at any extreme, its position reads as a coordinated solution to simultaneous constraints. Mammals spanning more than four orders of magnitude in brain volume^23^ converge on the same region, which suggests these constraints outweigh phylogenetic and volumetric diversity.

Within that pocket, brains are not centrally placed. They lean toward the capable, expensive edge of what is achievable, and so spend wiring to buy computation rather than minimising cost above all.

The rule that defines the space cannot itself build biologically completely similar networks. Even its best-fitting networks stay substantially farther from biological connectomes than from the reference and communication-optimised networks. Using a two-parameter fit to compress a connectome into the GNM morphospace, the following can be observed: The remaining error for biological networks is bigger than for the reference and communication-optimised networks, and the fitted parameters are not able to capture the changes in architecture that e.g. ages causes that the property space captures. The parameter-space is thus a more coarse embedding than the property space. Developmental age, for instance, should be recoverable from the fitted parameters^16^, yet here it is recovered far more clearly in the property space.

Maturation across development and divergence across evolution leave their strongest marks on the same structural dimension. In a property space fitted on the GNM networks alone, age moves the per-year consensus connectomes along a non-linear trajectory: first along PC2_GNM_ toward higher computational capacity, then, after a stable inflection near 16 years, along PC1_GNM_ toward greater integration. Higher PC1_GNM_ values go with higher wiring costs, so the first stage is consistent with the reported monotonic weakening of the wiring-length penalty across development^16^. PC2_GNM_ also carries the strongest phylogenetic signal, which may indicate a convergence between brain maturation and evolutionary processes.

Several limitations bound these conclusions. All connectomes derive from diffusion-MRI tractography, which is susceptible to crossing-fibre and false-positive artefacts^29^, so the results describe tractography-reconstructed networks rather than the ground-truth connectome. The property set is itself a choice, some of its measures rest on single stochastic estimates (one Louvain partition for modularity being the clearest case). Also, assigning properties to goals, architectural settings, and emergent behaviours is a deliberate organising choice whose groups are not perfectly orthogonal. Three dataset caveats apply on top. First, the HCP-D age labels may be uniformly offset by up to one year, which would shift the trajectory’s absolute position but not its shape or the inflection we report. Second, cortical folding varies with brain volume independently of taxonomic order^30^, so differences we attribute to phylogeny may partly or solely reflect volume-dependent geometry. Third, the two empirical datasets were acquired and reconstructed differently, in vivo against ex vivo and Schaefer-100 against endpoint clustering, so the offset between the HCP-D and MaMI clouds cannot be read as a biological difference.

Taken together, biological brain networks settle into a constrained, near-optimal compromise among cost, capability, and robustness. The compromise leans toward capability and robustness rather than cheapness. Development and phylogeny imprint on the same axis of that compromise (PC2_GNM_), so the variation separating a child from an adult runs parallel to the variation separating taxonomic orders. Testing that parallel directly can be done by utilizing a continuous evolutionary variable instead of taxonomic order, for example, the encephalisation quotient or divergence time from a common reference. This would place the developmental and evolutionary trajectories on one scale instead of comparing them by analogy. Furthermore, extending the pipeline to directed connectivity, to genetic regulatory gradients as generative priors, or to functional connectomes8 would provide further insights into how brain networks is shaped by maturational, genetic, and dynamical constraints.

## METHODS

### Empirical Datasets

#### Empirical Networks: Humans

Structural connectomes from 635 healthy individuals from the Human Connectome Project (HCP) Development dataset^18^ were used in the preprocessed form of Mousley et al.^31^, which applies the same preprocessing pipeline as^32^, and accessed in semi-processed format from DSI Studio’s Fiber Data Hub^33^. DSI Studio was used to reconstruct the structural connectomes with GQI-based deterministic tractography (parameters: maximum fibre length 250 mm, minimum fibre length 30 mm, 5 × 10^6^ streamlines, random seeding, step size 1 mm, and turning angle threshold 35*^◦^*). Parcellation was done in accordance to the Schaefer-100 atlas^34^. Count-pass streamline counting, in which a connection is registered for every region a streamline passes through, was used to determine connectivity. Then, the connectomes were binarised and thresholded to 10 percent density, using the function binarize_network from netneurotools^35^.

#### Empirical Networks Other Mammals

Structural connectomes from 224 mammals from the mammalian MRI (MaMI) dataset (acquisition^24^, reconstruction^19^, dataset^36^) were used. These connectomes span 12 taxonomic orders, from bats to giraffes, and were scanned ex vivo under a uniform protocol. Their acquisition, reconstruction (tractography with k-means parcellation), and distance harmonisation by the original authors are detailed in Supplementary Note S1.

Node count and density both affect graph measures, and so must be matched across datasets^37^. To match the 100-node human parcellation, each 200-node MaMI connectome was therefore downsampled to 100 nodes. Within each hemisphere, nodes were optimally paired by Euclidean distance (min_weight_matching, NetworkX ^38^) from a cdist distance matrix^39^. Each pair was then merged by averaging coordinates and summing connection weights, and the result was thresholded to 10% density and binarized (netneurotools^35^). MaMI distance matrices live on a different physical scale, so each was additionally *z*-score-rescaled to the Schaefer-100 distance distribution,

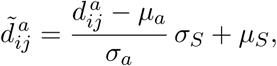

where *µ_a_, σ_a_* are the mean and standard deviation of animal *a*’s distances and *µ_S_, σ_S_* those of the Schaefer-100 distribution. This preserves relative structure while matching the human mean and standard deviation for cross-species wiring-cost comparison. One animal - an African green monkey that remained disconnected after downsampling - was removed. Order-level analyses used the five orders with *n >* 20 (Primates, Carnivora, Cetartiodactyla, Rodentia, and Chiroptera).

#### Age-stratified Consensus Networks

For the developmental analysis, individual HCP-D connectomes were grouped by integer age and a consensus network was computed per age with struct_consensus (netneurotools), then thresholded to 10% density and binarized. This yielded 17 consensus networks spanning ages 6-22.

### Synthetic Datasets

The concept of the *morphospace* originates in theoretical biology, where it maps the space of possible phenotypes^20^. For networks, the axes can be defined in two ways^12^. They can either be network properties, so that each point is a measured architecture (e.g. routing against diffusion efficiency). Alternatively, they can be the parameters of a generative model, so that each point is a distinct wiring rule. Both have been used for brain networks: the first to map functional organisation^21^, the second to map networks by a small set of topological properties^12^. This work uses both. The property space is built from what the networks *are*, its axes being principal components of the 29 measured properties, whereas the GNM morphospace is built from how they were *made*, its coordinates being the generative parameters *η* and *γ*. Each connectome therefore holds two positions: one set by its measured properties, and one by the wiring rule that comes closest to reproducing it.

#### GNM Morphospace: Generative Network Model (GNM)

A generative network model builds a network from a rule rather than from data^15^. It starts from a sparse seed and adds one edge at a time until a target density is reached. Which edge is added is decided by a probability that trades two competing pressures against each other. The first is the metabolic cost of the connection, approximated by the distance it spans. The second is its topological value, approximated by how far the two regions’ existing neighbourhoods already overlap (homophily). Two exponents, *η* and *γ*, set how heavily each pressure counts, so one (*η, γ*) pair fully specifies one wiring rule. Sweeping them produces a controlled family of architectures, each the outcome of a different cost-versus-value compromise. That is what makes the model a source of possible architectures rather than a fit to one particular brain.

At each growth step the GNM assigns a connection probability to each candidate node pair (*i, j*) according to:

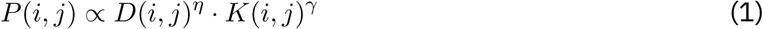

where *D*(*i, j*) is the Euclidean distance between nodes *i* and *j* in the Schaefer-100 parcellation and *K*(*i, j*) is the matching index^16^, a normalised measure of how far the two nodes’ neighbourhoods overlap. The two exponents pull in opposite directions. Negative *η* makes a long connection unlikely, and so prices distance; the more negative it is, the harsher the price. Positive *γ* makes a connection between two similarly connected regions more likely, and so rewards homophily; the larger it is, the stronger the reward. Setting *η >* 0 inverts the distance term and rewards long connections instead, which is metabolically implausible for a brain. We include that regime so that the sweep brackets the biologically plausible one rather than assuming it. The matching index was selected as the homophily measure because it produces the closest fit to empirical connectomes among competing homophily rules^22^. All networks are binary and undirected (i.e. the adjacency matrix *A* is defined as *A_ij_* ∈ {0, 1}), with node coordinates fixed to the Schaefer-100 atlas so that *D*(*i, j*) reflects realistic cortical geometry. The growth process is seeded from a minimum spanning tree (MST) computed on the Schaefer-100 distance matrix via minimum_spanning_tree from NetworkX ^38^, to ensure full connectivity across the (*η, γ*) parameter space. Edges are added one at a time until the network reaches the empirical target density, with *K*(*i, j*) recomputed after each addition. The parameter space was covered by a 50 × 50 grid with *η* ∈ [−8, 3] and *γ* ∈ [−0.1, 1]; 10 replicate networks were drawn per grid cell, yielding 25,000 synthetic networks in total. The GNM implementation is from^40^.

#### Erdős-Rényi (ER) Random Networks

As a structural baseline, 100 Erdős-Rényi random networks^41^ were generated on *n* = 100 nodes with fixed edge density 10 %. Each network was generated by sampling exactly *m* = 495 edges uniformly at random without replacement from all 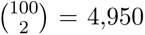 possible node pairs. This yielded binary, undirected adjacency matrices with identical density across all instances. The Schaefer-100 Euclidean distance matrix used for the GNM was passed to spatially sensitive properties on the ER ensemble.

#### Minimum-Wiring Network

All 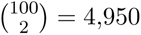 possible node pairs were ranked by Euclidean distance from the Schaefer-100 brain coordinate matrix, and the *m* = 495 shortest-distance pairs were connected (density 10 %) - yielding one deterministic, binary, undirected adjacency matrix. Every node has at least four connections, and the network forms a single connected component.

#### Communication-Optimised Reference Networks

The GNM sweep spans architectures built from a cost-versus-homophily compromise. It says nothing about what an architecture looks like when one particular job is done as well as possible. The communication-optimised reference networks fill that gap. Each is grown to be as good as it can be at one specific model of how a signal travels through a network, with no other objective allowed to interfere. They are therefore interpretable landmarks, not null models.

The three models differ in how much a signal is assumed to know about the network^10^. *Routing* assumes the signal takes the shortest path, as if it knew the whole topology in advance; optimising it therefore favours short characteristic path lengths and a few long-range shortcuts. *Diffusion* assumes the opposite: the signal has no knowledge and wanders as an unbiased random walk, so optimising it favours architectures in which a random walker reaches any target quickly. *Propagation* sits between the two - the signal is broadcast along all paths simultaneously, with longer paths discounted - so optimising it favours networks that are richly interconnected at every scale.

To construct them, we utilised a game-theoretic generative framework described in^42^ and implemented in the open-source Python library YANAT ^43^. Briefly, this framework constructs networks by iteratively adding edges that maximise a target communication objective, evaluated via Shapley value decomposition of nodal contributions. The three objectives are global efficiency (routing), random-walk commute time (diffusion), and communicability (propagation). All optimised networks were seeded with a ring lattice and grown to a target density of 10%, matching the density of our empirical and synthetic connectomes. Our target density differs from the 20% used in^42^. The resulting trade-off parameters (*α*) are therefore not directly comparable to those reported there (see Supplementary Table S1).

#### Properties

The 29 properties were first used to analyse the property space without any grouping of the properties. Only later, each property was manually assigned into a structure of properties (see Fig. 1E).

The 29 properties were chosen to be non-redundant as possible.

The following will present first the structure of properties. Then, one representative of each group is described in more detail. Every property’s values over the 25,000 GNM networks are visualised in Supplementary Fig. ??, and the remaining information on all properties can be found in Supplementary Note 2.

Structure of the Properties

The 29 properties are organised at three levels (Figure 1E). At the top are three *goals*, on which selection acts directly because they are assumed to be the main drivers of fitness: economy (be cheap, i.e. minimise wiring and signalling cost), capability (be able, i.e. memory and dynamical flexibility), and robustness (be durable). Each goal is measured by outcome properties quantifying how much of it was achieved (e.g. memory capacity reads out capability; targeted-attack AUC reads out robustness). The goals are realized by *architectural* features. We divide the features into three groups, each as independent of the others as possible: reach (how far edges go), centralization (how concentrated connectivity is on a few hubs), and segregation (how modular the network is). Because a single architectural setting affects several goals at once, the groups are necessarily non-orthogonal. Finally, *emergent behaviour* - communication (signalling: routing, diffusion, broadcast) and near-criticality (the dynamical regime, captured by the spectral properties) - are emergent properties that arise from the architecture and, in turn, feed into the goals.

#### Representative Properties

One property per group is summarised (very briefly) here; Figures 7 and 8 show the four experiments behind the three goal readouts. One full description of a representative property, with its parameters and formulas, is given in Supplementary Note 2, and all 29 properties are defined in Supplementary Note S1.

**Figure 8:**
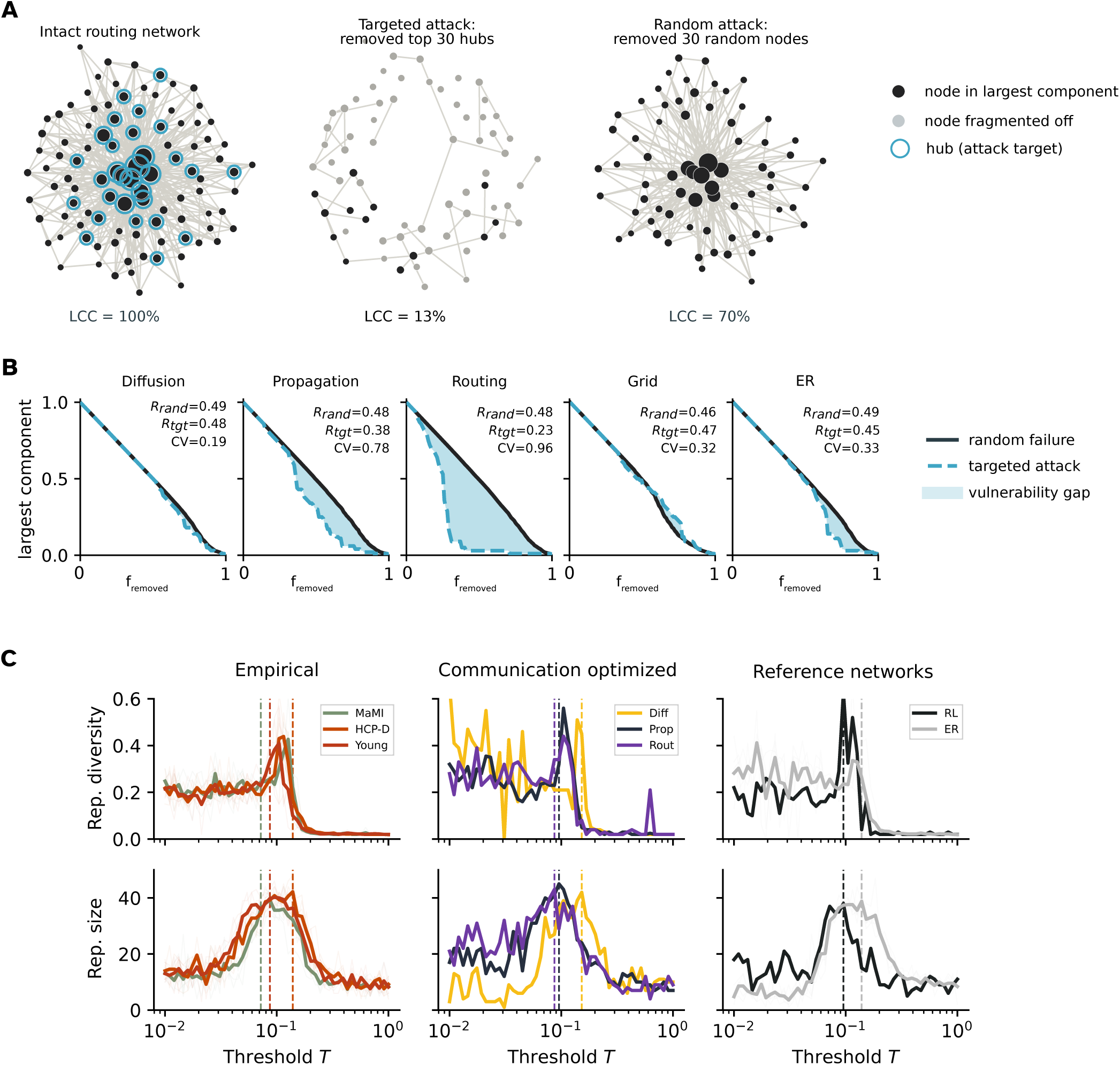
The experiments behind the robustness and critical-dynamics readouts. (A) Attack tolerance illustrated on the routing-optimised network: intact, after removal of its 30 highest-degree hubs, and after removal of 30 random nodes. Black, nodes in the largest connected component; grey, nodes fragmented off; blue rings, the targeted hubs. LCC gives the size of the largest component as a percentage of all nodes. (B) Largest component against the fraction of nodes removed, under random failure (black) and targeted attack (blue dashed), for five reference networks. Shading marks the vulnerability gap between the two curves; annotations give both AUCs (*R*_rand_, *R*_tgt_) and the degree heterogeneity CV. (C) Critical-dynamics sweep, averaged over 20 runs per network: repertoire diversity (top) and repertoire size (bottom) against the activation threshold *T* , for empirical connectomes, communication-optimised networks, and structural baselines (left to right). The critical threshold is indicated by the dashed vertical lines; fine lines represent individual networks, bold lines represent the average across datasets.

*Economy* is read out as the fraction of long-range connections *f*_LR_, the proportion of edges longer than the 39.56th percentile of the empirical edge-length distribution, which is where the 9 cm long-range cut-off of^28^ falls. At this threshold *f*_LR_ tracks absolute wiring cost *W* =Σ*ij a_ij_d_ij_* ^5^ at *r* = 0.997 across the 25,000 synthetic networks, the strongest of four candidate percentiles (Figure 7A), so it serves as a parcellation-invariant proxy for cost. Economy is reported as the complement 1 − *f*_LR_, so that higher values mean cheaper wiring.

*Capability* is read out twice, because its memory and its dynamical aspects need different simulations. For memory, each connectome served as the recurrent weight matrix of an echo state network44 implemented in echoes17, spectrally normalised to *ρ* = 0.9 and run with leakage *α* = 0.3 and input scaling *s* = 0.1. Capacity was decomposed with the information processing capacity framework^45^; its degree-1 term IPC_1_, equivalent to linear memory capacity, is the property we retain, averaged over 20 trials (Figure 7B, C). For dynamics, an excitable cellular-automaton model^46^ was run on the distance-weighted, spectrally normalised connectome across a logarithmic sweep of the activation threshold *T* . The threshold at which the repertoire of activity patterns peaks defines the critical threshold *T_c_*, and the repertoire size and diversity at *T_c_* are retained (Figure 8C).

*Robustness* is read out by removing nodes one at a time and tracking the largest connected component, either in random order or in decreasing order of degree, recalculated after each removal^9^. The area under the resulting curve (AUC) quantifies resilience. Because the two regimes dissociate along degree heterogeneity (Figure 8A, B), we retain the targeted-attack AUC as the primary robustness property and additionally report the targeted-to-random ratio, which isolates hub-driven vulnerability from overall connectedness.

The architectural settings and emergent behaviours are measured by quite established graph statistics and are therefore only named here. *Reach* uses Ollivier-Ricci curvature^47^ on an *α*-lazy random walk (*α* = 0.5), computed with GraphRicciCurvature^48^ and summarised per network by its minimum and skewness. *Centralization* uses the standard deviation across nodes of average controllability, derived from the controllability Gramian of a linear time-invariant model of the connectome^49,50^. *Segregation* uses modularity *Q* ^51^ over a Louvain partition^52^. *Communication* uses global efficiency, the mean inverse shortest-path length^53^. *Near-criticality* uses the spectral radius *ρ*(*A*) = max*_i_* |*λ_i_*(*A*)|, which separates activity that decays (*ρ <* 1) from activity that amplifies (*ρ >* 1)^6^. The last three use the NetworkX implementations^38^.

### Property space construction via PCA

To generate the property space (see Section 2.1), we assembled the property matrix spanning all networks: 25,000 GNM networks, HCP-D connectomes (*n* = 635), MaMI connectomes (*n* = 224), structural baselines (*n* = 101), and communication-optimised reference networks (*n* = 3), for a total of 25,963 networks. Each property was then *z*-scored independently across these 25,963 networks (zero mean, unit variance), and the standardised matrix was submitted to principal component analysis (PCA) using the decomposition.PCA implementation in scikit-learn^54^. The resulting principal components define the property space in which all networks are jointly positioned. Property loadings (Figure 4).

### Pairwise and Ternary Trade-Off Analyses

Pairwise trade-off plots (Figure 3A-B) show one property against another across all networks, so that the visible Pareto front reflects the true achievable boundary. The ternary (triangle) plot (Figure 3C) summarises a network’s *relative* balance across three properties - economic efficiency (1 − *f*_LR_), global efficiency, and memory capacity (here the standalone forgetting-curve estimate at input scaling *s* = 0.1, not the IPC_1_ used in the PCA; see above). The per-axis mapping is rank (quantile) normalisation. For each property, a network’s raw value is replaced by its empirical-CDF position, that is its rank divided by the sample size, within the pooled distribution of all plotted networks. Each axis thereby becomes uniform on [0, 1] regardless of its original scale or skew, which spreads the cloud across the full simplex so that all three vertices and edges are reachable. A network’s coordinates then encode its relative standing on each property rather than an absolute value. It approaches a corner when it ranks high on one property and below the median on the other two, and sits near the centre when the three ranks are balanced. The projection therefore reveals how a network’s three properties compare to one another, but not their common level: a network ranking high on all three and one ranking low on all three both map to the centre. To recover that level, we additionally render the same three rank-normalised axes as a triangular prism (Supplementary Fig. S5). Each network keeps its ternary floor position and is lifted to a height equal to the sum of its three rank scores (range [0, 3]), so the vertical axis measures a network’s *total* standing across economic efficiency, global efficiency, and memory capacity at once, while the floor still shows their relative balance.

### Developmental Trajectory and Phylogenetic Analyses

To ask whether biological variation aligns with the model’s own axes of variation - rather than axes partly defined by the biological data - a separate PCA was fit exclusively on the 25,000 GNM networks, restricted to the 29 selected properties, defining the GNM-fitted property space. We refer to the axes of this GNM-fitted space as PC1_GNM_ and PC2_GNM_ to distinguish them from the joint property space axes. This PCA was computed on the properties in their non-inverted (raw) direction. Per-year HCP-D consensus connectomes, yielding 17 observations spanning ages 6-22 years, and individual MaMI connectomes were then projected into this space using the scaler and PCA transformation fit on the GNM data alone. Smoothing splines were fit independently to the PC1_GNM_ and PC2_GNM_ coordinates of the consensus observations as a function of age using scipy.interpolate.make_smoothing_spline^39^ with smoothing parameter *λ* = 10, evaluated at 300 uniformly spaced time points. The choice *λ* = 10 lies within a stable regime: as shown in Supplementary Note S8, the identified inflection age is stable at approximately 16 years for all *λ* ≥ 5.

The inflection age, i.e. the age of maximum trajectory curvature, was localised by computing the curvature at each time point as

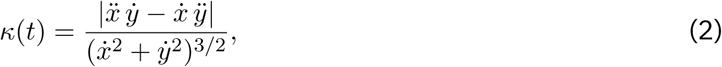

where *x* and *y* are the smooth PC1_GNM_ and PC2_GNM_ curves as a function of age, and dots denote derivatives with respect to age. The age at which *κ*(*t*) is maximal was taken as the developmental inflection point.

Pearson correlations were computed between age and each of the first ten PC_GNM_ scores across the 17 per-year consensus observations. Permutational multivariate analysis of variance (PERMANOVA)^25^ pseudo-*F* statistics were computed per taxonomic order (one-vs-rest) for each of the first ten PC_GNM_ axes across individual MaMI connectomes; the mean pseudo-*F* across orders was used to rank components by phylogenetic discriminability.

We tested pairwise separation between the five MaMI orders with *n >* 20 in the PC1_GNM_-PC2_GNM_ plane. The cut at *n >* 20 is clean: the largest excluded order holds *n* = 9 individuals, and the smallest included one *n* = 29. For each pair we combined a permutation-based significance test with a multivariate effect size. PERMANOVA^25^ (implemented in scikit-bio^55^) compares the average between-group to within-group Euclidean distance via a pseudo-*F* statistic, and derives a *p*-value by permuting group labels (999 permutations). It therefore makes no distributional assumption and tolerates the unequal group sizes of the MaMI dataset, which classical MANOVA (covariance homogeneity, multivariate normality) and per-PC ANOVA (multiple testing, single axis only) do not. Because significance scales with sample size, we additionally quantified the magnitude of each separation as the Mahalanobis distance between group centroids,

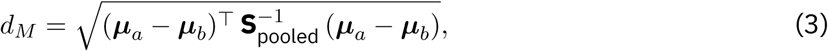

where ***µ****a,* ***µ****b* are the group centroids of order *a* and *b*, and S_pooled_ is the pooled within-group covariance matrix, that is the within-group covariances of the two orders averaged together and weighted by their sample sizes to give one shared estimate of within-group spread. Normalising by S*−*1_pooled_ rescales each axis by its variance, so that the higher-variance PC1_GNM_ and the lower-variance PC2_GNM_ contribute comparably. *d_M_* is the multivariate analogue of Cohen’s *d*, the standard effect size expressing a difference of group means in units of within-group standard deviation, and is read on the same scale (*d_M_* ≼ 0.3 negligible, *d_M_* ≈ 1 one pooled standard deviation). The two measures answer complementary questions: PERMANOVA tests whether a separation is statistically real, and *d_M_* how large it is relative to within-group spread. Together they distinguish biologically meaningful differences from merely detectable ones.

### Comparing Networks: DeltaCon

To evaluate similarity between networks and to embed them in the GNM morphospace, we used *DeltaCon*^27^. We chose it over the field’s KS-based Energy distance^16^ on the evidence of a companion benchmark, in which we scored 16 candidate distance measures^56^. DeltaCon’s decisive advantage there is speed combined with high accuracy: for instance, comparing all HCP-D connectomes to all 25,000 synthetic networks took approximately 23 CPU-core hours instead of slightly more than 13 CPU-core days, which the field’s default KS-based energy would have required.

DeltaCon computes node-affinity matrices *S* via fast belief propagation,

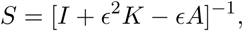

where *I* is the identity matrix, *K* the diagonal degree matrix, *A* the adjacency matrix, and *c* a small constant governing neighbour influence decay. The affinity matrix *S* captures influence flow across all path lengths, with indirect paths contributing progressively less. The distance between two graphs *G* and *G^t^* is then computed as the Matusita distance^57^ between their respective affinity matrices:

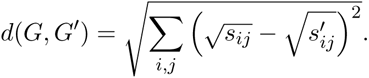

Unlike simple edge-counting measures, DeltaCon is sensitive to the structural role of connections - for instance, removing a bridge edge in a barbell graph produces a much larger distance than removing a redundant intra-clique edge, reflecting the disproportionate impact on information flow. DeltaCon is independent of the lengths of the connections. We used the implementation from netrd^58^.

Each connectome was assigned the (*η, γ*) of the single most similar GNM network (lowest DeltaCon) as its *best-fit GNM morphospace position* (dots in Figure 6; input to the PERMANOVA of Supplementary Fig. S8 and Supplementary Table S5). Because the minimum is taken over individual networks, a dot need not coincide with the lightest heatmap cell. To summarise a dataset by a single position, we took the *medoid* of its best-fit coordinates, that is the coordinate minimising the summed Euclidean distance, in raw (*η, γ*) units, to every other best-fit coordinate in that dataset. Unlike a mean, the medoid is an actual member of the set, so it always corresponds to a GNM that was generated. Separately, averaging DeltaCon across the 10 realisations of each (*η, γ*) cell gives a mean-distance surface. The mean over that surface’s 100 lowest cells defines each dataset’s *residual distance*, that is how close a GNM can *reliably* get to a network, rather than the minimum distance one good-fitting network may reach by chance.

### Statistics and Reproducibility

All statistical analyses were performed in Python. Sample sizes were not predetermined by a power analysis; we analysed all available connectomes in each dataset (HCP-D: *n* = 635; MaMI: *n* = 224 after excluding one disconnected animal, Section 4.1) alongside 25,000 GNM networks, 100 Erdős-Rényi networks, one minimum-wiring network, and three communication-optimised reference networks. Beyond the single excluded MaMI connectome noted above, no data were excluded, and no blinding or experimental randomisation applies to this observational, computational study. Age-related change was assessed with two-tailed Pearson correlations between age and each of the first ten PC_GNM_ scores across the 17 per-year HCP-D consensus observations; a correlation was considered significant at *p <* 0.05. Differences between mammalian orders were tested with PERMANOVA^25^ (scikit-bio^55^) using Euclidean distances and 999 label permutations, restricted to the five orders with *n >* 20 (Carnivora, *n* = 51; Primates, *n* = 46; Cetartiodactyla, *n* = 40; Rodentia, *n* = 29; Chiroptera, *n* = 29); effect sizes were quantified as the Mahalanobis distance between order centroids (Section 4.6). Reported pseudo-*F* , *r*, and *p* values are given in the Results and figure legends. Stochastic computations were averaged over repeated runs as stated for each property (memory capacity: 20 trials; critical dynamics: 20 runs per network; GNM: 10 replicates per (*η, γ*) grid cell).

## DATA AVAILABILITY

The mammalian MRI (MaMI) connectome dataset analysed in this study is publicly available on Zenodo under a CC BY 4.0 licence^36^; its acquisition is reported in^24^ and its reconstruction in^19^. The Human Connectome Project Development (HCP-D) diffusion data are available from the Human Connectome Project^18^, subject to its data use terms, and were accessed in semi-processed form through the DSI Studio Fiber Data Hub^33^. The networks generated for this study, namely the 25,000 synthetic GNM networks and the reference networks, alongside the computed property matrices that underlie all figures, will be made available on Zenodo at the time of publication. That deposit will also redistribute our preprocessed 100-node MaMI connectomes and their distance matrices, which are a modified version of the CC BY 4.0 MaMI data^36^; the modifications are those described in Section 4.1 and are itemised in the deposit. HCP-D connectomes are not redistributed in any form, in keeping with the Human Connectome Project data use terms.

## CODE AVAILABILITY

All custom analysis code is implemented in Python and will be made openly available at the time of publication.

The analysis relies on the open-source libraries NetworkX 38, generativenetworkmodels40, netneurotools35, echoes59, YANAT 43, the Brain Connectivity Toolbox for Python (bctpy)60, GraphRicciCurvature48, netrd58, scikit-learn54, scikit-bio55, SciPy39, and NumPy 61.

A large language model (Anthropic Claude, via Claude Code) was used during this project.

## ACKNOWLEDGEMENTS

A.D. was supported by the Max Weber Programme of the Elite Network of Bavaria and by the Erasmus+ programme of the European Union. K.F. and D.E.A. were supported by the Templeton World Charity Foundation, Inc. under grant TWCF-2022-30510. F.P.d.S. was funded through the EU project Virtual Brain Twin (Horizon Europe; No 101137289). D.E.A. was also supported by the James S. McDonnell Foundation Opportunity Award and by a Mental Health Award from the Wellcome Trust (309245/Z/24/Z). All research in the Department of Psychiatry at the University of Cambridge is supported by the National Institute for Health and Care Research Cambridge Biomedical Research Centre (NIHR203312) and the NIHR Applied Research Collaboration East of England.

## AUTHOR CONTRIBUTIONS

A.D., K.F., and D.E.A. conceived and designed the study. A.M. contributed the HCP-D connectome processing on which the human data are based. F.P.d.S. contributed to the implementation of the dynamical-repertoire properties. A.D. and K.F. developed the analysis pipeline, generated the synthetic and reference networks, performed all computations and statistical analyses, prepared the figures, and wrote the manuscript. K.F. and D.E.A. supervised the project and contributed to the interpretation of the results and to editing the manuscript. All authors read and approved the final manuscript.

## COMPETING INTERESTS

The authors declare no competing interests.

## MATERIALS & CORRESPONDENCE

Correspondence and requests for materials should be addressed to A.D.

## Notes

https://zenodo.org/records/6376544

### Competing Interest Statement

The authors have declared no competing interest.

