## Supplemental Material for "Mapping the Trade-Offs That Shape Mammalian Brain Network Architecture"

Supplementary figures and tables are numbered S1, S2, and so on. Unprefixed references to "Fig.", "Table", "Section", or "Methods" point to the main article. The sections below cover the mammalian dataset, the communication-optimised reference networks, the full set of 29 network properties and their landscapes across the synthetic networks, how the two morphospaces are built and read, additional property-space visualisations, further trade-off analyses, and finally the developmental, phylogenetic, and distance-measure analyses.

### DATASETS

#### Mammalian Dataset

The MaMI connectomes were acquired and reconstructed by their original authors, our further processing is described in Section 4.1. The 225 connectomes, one of which we exclude, cover 124 species across 12 taxonomic orders. Acquisition is described in [1], reconstruction in [2], and the original dataset itself is available on Zenodo [3]. Brains were collected after natural or incidental death, then extracted and fixated within 24 hours. The scanning was performed on one of two scanners, chosen by brain size, and acquired both high-resolution anatomical scans (T2- or T1-weighted MRI) and diffusion MRI (high angular resolution diffusion imaging). Then, diffusion analysis and tractography followed, after which "certain fiber tracts were removed" [2].

Parcellation was then obtained by k-means clustering of the fibre end-point positions within each hemisphere separately, into 200 nodes whose locations are the cluster mass centres; each connection weight is the number of streamlines between the two nodes. Because the cluster count is fixed, this parcellates brains of any species into the same number of regions [2]. Alongside the node locations and connection matrices, the dataset provides the taxonomic label of each animal. Then, we continued processing the data as described in Section 4.1.

#### Communication-Optimised Reference Networks

The communication-optimised reference networks were grown to a target density of 10 %, whereas the original work [4] used 20 %. Because the trade-off parameter $\alpha$ balances the communication objective against wiring cost at a given density, the values obtained here are not directly comparable to the published ones (see Table S1).

| Type | Schaefer 100 (here) | Schaefer 100 ([4]) |
|---|---|---|
| Routing | 13.16 | 43.01 |
| Diffusion | 69.12 | 8.13 |
| Propagation | 3.65 | 10.46 |

Table S1: **Trade-off parameters ($\alpha$) of the communication-optimised reference networks.** The $\alpha$ values that resulted for the networks we created here (10±1 percent density) in comparison to the $\alpha$'s utilised in[4] (20±1 percent density).

### PROPERTIES

In the following, all 29 properties are described in two passes: This section takes one representative property per group and describes the method of computing it in full. Capability is illustrated twice, by its memory readout and by its dynamical readout, because the two are measured by different simulations. Section S1 then tabulates all 29properties, with each properties's source, reasoning why it is useful, and formula, and maps each one across the $\eta$-$\gamma$ landscape of the $25{,}000$ synthetic networks. Both passes follow the three-level scheme used throughout this work: the three goals, the three architectural settings, and the two emergent behaviours (Section 4.3).

**Economy: Fraction of long-range connections** The fraction of long-range connections $f_{\text{LR}}$ is the proportion of a network's edges that count as "long-range". We first defined the cut-off from which we describe a connection as long range: For that, we took the cut-off from[5], who call connections longer than 9 cm long-range - however, since HCP-D and MaMI use different scalings and include differently large brains, this absolute threshold is not suitable for both datasets. Thus, we determined that 39.56% of the connections in the HCP-D networks exceed 9 cm, and then placed the threshold for either dataset at the 39.56th percentile of the empirical edge-length distribution to make the human and the mammalian dataset comparable. We then verified that, defined this way, $f_{\text{LR}}$ is a parcellation-invariant proxy for absolute wiring cost $W = \sum_{ij} a_{ij} d_{ij}$[6]: across all 25,000 synthetic networks the two correlate at $r = 0.997$, the strongest among four candidate percentiles (10%, 30%, 39.56%, 50%; Fig. 7A). Expressing wiring cost as a fraction of long-range connections is convenient because it accounts for the generally different distance-distribution curves between datasets.

Wiring economy is reported as the complement $1 - f_{\text{LR}}$, so that higher values reflect cheaper, more local wiring.

A note on the wiring cost property and the distance-weighted dynamical measures: To enable a cross-species analyses, MaMI distance matrices were rescaled to match the mean and standard deviation of the Schaefer-100 atlas distance distribution before the threshold was computed.

**Capability: Memory capacity (linear)** Memory capacity was assessed with echo state networks (ESNs)[7,8] as implemented in the `echoes` Python library[9]. The structural connectivity matrix served as the reservoir's recurrent weight matrix $W_{\text{res}}$, and was spectrally normalised to $\rho = 0.9$. Reservoir dynamics then followed the leaky-integrator update:

$$\mathbf{x}(t+1) = (1-\alpha)\,\mathbf{x}(t) + \alpha\,\tanh\big(W_{\text{res}}\,\mathbf{x}(t) + W_{\text{in}}\,u(t)\big), \tag{S1}$$

with leakage rate $\alpha = 0.3$ and input scaling $s = 0.1$. We quantified what each reservoir can compute with the information processing capacity (IPC) framework[9,10], which decomposes the total capacity of a reservoir into contributions from orthogonal target functions of increasing polynomial degree. The reservoir received a scalar i.i.d. input $u(t)$ and was trained via ridge regression to reconstruct degree-$d$ polynomial functions of delayed inputs $u(t-\tau)$ (delays $\tau \in \{1,\ldots,40\}$)[10,11]. Then, the capacity for a single target $y_k$ was defined as $\text{IPC}_k = (P\,y_k)^2/\|y_k\|^2$, where $P$ is the projection onto the reservoir-state space, and the degree-$d$ capacity $\text{IPC}_{\text{deg}_d} = \sum_k \text{IPC}_k$ sums these over all degree-$d$ targets. In this work, we only look at $d = 1$ and $d = 2$. The degree-1 (linear) capacity $\text{IPC}_1$ quantifies delayed recall of the input signal and is therefore equivalent to the linear memory capacity. Higher values reflect more temporally extended information storage. The degree-2 (nonlinear) capacity $\text{IPC}_2$ quantifies recall of squared delayed inputs, and therefore describes how well the network can compute non-linear functions of its remembered inputs. Results were averaged across 20 independent trials per network (each with a different random input projection $W_{\text{in}}$)[9]. Mean linear and non-linear capacity per dataset, and the decay of linear capacity across input delays, are shown in Figure 7.

Two figures use a second, self-implemented memory capacity estimator, which returns memory capacity on a per-lag basis: Figure 7, which plots capacity against lag, and the memory-capacity axis of the ternary trade-off triangle (Figure 3C, and its prism rendering in Figure S5). Figure 7 requires this second estimator, because its purpose is to show *how* computational capacity is measured and how it decays with input delay, which the summary $\text{IPC}_1$ cannot display. The ternary triangle does not require it, and the use of the second estimator in that panel is mostly a historical artifact without much influence, as the figure only utilizes the *relative* rank ordering of networks by memory capacity.

The two estimators agree due to their setup, as both the degree-1 IPC and the linear memory capacity from the second estimator fit a linear readout on training reservoir states and score the squared correlation on held-out data. The figures are therefore a faithful conceptual depiction of the measure but not a literal, one-to-one rendering of the $\text{IPC}_1$ values used in the analysis. The routine for the second estimator averages over 500 trials and obtains the readout from the `echoes` solver directly, whereas the analysis $\text{IPC}_1$ averages over 20 trials and fits the readout via an explicit pseudo-inverse of the reservoir states.

**Capability: Critical dynamics.** Beyond memory, the activity dynamics are also of interest to evaluate capability. Here, critical dynamics were defined as the richness of the activity-pattern repertoire generated by an excitable cellular-automaton model on the structural connectome[12]. The connectivity matrix was first weighted by the inverse inter-regional distance ($\tilde{A}_{ij} = A_{ij}/d_{ij}$) and then spectrally normalised to a spectral radius of $1$, so that the dynamics are comparable across networks. Each node occupies one of three states - quiescent ($Q$), excited ($E$), or refractory ($R$). At each step a quiescent node becomes excited if the weighted input from its excited neighbours exceeds the activation threshold $T$ (or spontaneously, with a small noise probability); an excited node deterministically becomes refractory; and a refractory node returns to quiescence with a fixed probability[12]. For each network the model was simulated for $1{,}000$ steps at every threshold on a logarithmic sweep $T \in [10^{-2}, 1]$, and the repertoire size (the number of unique network-wide activity patterns visited) and repertoire diversity (their heterogeneity) were recorded. The critical threshold $T_c$ was defined as the value of $T$ at which the repertoire size peaks; the repertoire size and diversity at $T_c$ were retained as the critical-dynamics properties, quantifying how rich a range of activity patterns the network supports near its dynamical transition. The per-dataset repertoire sweeps are shown in Fig. 8C.

**Robustness: Random and targeted attacks** Robustness (see e.g.[13,14]) was assessed by simulating sequential node removal and tracking the size of the largest connected component (LCC) as a fraction of the original network size. Two attack strategies were evaluated: *random attack* removes nodes in uniformly random order, simulating non-specific failure; *targeted attack* removes nodes in decreasing order of degree at each step (recalculated after each removal) and thus mimics disruption of the most central nodes[13]. Robustness is quantified as the area under the LCC curve (AUC) as a function of the fraction of nodes removed, normalised to $[0, 1]$. Higher AUC indicates that the LCC degrades slower and the network is more resilient. For both strategies, removal was simulated until all nodes were removed, and AUC was computed.

The two regimes dissociate along degree heterogeneity: in homogeneous networks every node contributes comparably, so the two AUCs nearly coincide. In contrast, in hub-dominated, heavy-tailed networks a few high-degree nodes hold the topology together, which makes them resilient to random failure but fragile once those hubs are removed first. This is the classic structural cost of hub-mediated integration (Figure 8)[13]. We retain the targeted-attack AUC as the primary robustness property and additionally report the ratio of targeted-to-random AUC, which isolates this hub-driven vulnerability from a network's overall connectedness.

**Reach: Ollivier-Ricci curvature** Ollivier-Ricci curvature compares the transport cost to geodesic distance[15]. For an edge $(x, y)$ it is here defined as

$$\kappa(x, y) = 1 - \frac{W_1(\mu_x^\alpha, \mu_y^\alpha)}{d(x, y)}, \tag{S2}$$

where $\kappa(x, y)$ is the curvature of the edge, $d(x, y)$ is the shortest-path distance between its endpoints, and $W_1$ is the Wasserstein-1 (earth-mover's) distance between the two $\alpha$-lazy random-walk distributions defined as

$$\mu_x^\alpha = (1 - \alpha)\, \delta_x + \alpha\, \text{Uniform}(N(x)). \tag{S3}$$

Here $N(x)$ is the set of neighbours of node $x$, $\text{Uniform}(N(x))$ places equal mass $1/|N(x)|$ on each of the neighbours of x, $\delta_x$ is the point mass at $x$ itself, and the laziness parameter $\alpha \in [0, 1]$ sets how much mass leaves $x$: the walker stays put with probability $1 - \alpha$ and steps to a uniformly chosen neighbour with probability $\alpha$. We use $\alpha = 0.5$. An edge has positive curvature when the neighbourhoods of its endpoints are easy to transport into one another, i.e. they share many parallel paths (a redundant, well-clustered region). An edge has negative curvature when transport is costly because the edge is a bottleneck or bridge whose removal lengthens local routes. Curvature values near zero indicate a tree-like structure. Curvature is thus a local, mesoscale geometric descriptor of reach that global path measures might miss. The distribution of edge curvatures has been shown to characterise non-homogeneous network topologies[16] and to discriminate robust from fragile regions in brain structural connectivity[17]. We compute $\kappa$ for every edge with the `GraphRicciCurvature` library[18], that is based on[16], and summarise the resulting distribution per network by its minimum, and skewness.

**Centralization: NCT controllability standard deviation** Network control theory (NCT) models the structural connectome as the adjacency matrix of a linear time-invariant dynamical system[19,20]:

$$\mathbf{x}(t+1) = A\,\mathbf{x}(t) + B\,\mathbf{u}(t), \tag{S4}$$

where $\mathbf{x}(t)$ is the brain state vector, $A$ is the normalised adjacency matrix, $B$ selects a set of control nodes, and $\mathbf{u}(t)$ is the control input. Average controllability of a node is defined as the trace of the controllability Gramian $W_K = \sum_{t=0}^{\infty} A^t BB^\top (A^\top)^t$ when that node alone is the controller[19]: this quantifies the ease of steering the network towards all reachable brain states from that node on average. The NCT controllability standard deviation (NCT-STD) is computed as the standard deviation of per-node average controllability values across all $n$ nodes. A high NCT-STD indicates a heterogeneous network in which control capacity is concentrated in a few high-degree hub regions; a low NCT-STD indicates homogeneous, evenly distributed control capacity[19]. The adjacency matrix was normalised by dividing by $\lambda_{\text{max}}(A)+1$ prior to Gramian computation to ensure stable dynamics[20].

**Segregation: Modularity.** Modularity $Q$[21] quantifies the degree to which a network can be partitioned into internally dense and externally sparse modules. It can be calculated as

$$Q = \frac{1}{2m}\sum_{ij}\left[A_{ij} - \frac{k_i k_j}{2m}\right]\delta(c_i, c_j), \tag{S5}$$

where $A_{ij}$ is the adjacency matrix, $k_i$ is the degree of node $i$, $m$ is the number of edges, and $\delta(c_i, c_j) = 1$ if nodes $i$ and $j$ belong to the same community. Community assignments were obtained using the Louvain algorithm[22] as implemented in `NetworkX` (`community.louvain_communities`); modularity was then computed via `community.modularity()` based on the work of[23].

**Communication: Global efficiency** Global efficiency[24,25] is defined as the average inverse shortest-path length over all node pairs:

$$E_{\text{glob}} = \frac{1}{n(n-1)}\sum_{i\neq j}\frac{1}{spl_{ij}}, \tag{S6}$$

where $spl_{ij}$ is the shortest-path length between nodes $i$ and $j$ in the binary graph (set to $\infty$ for disconnected pairs, contributing zero to the sum). It captures the efficiency of parallel information transfer across the whole network and is inversely related to the characteristic path length; values near 1 indicate a densely interconnected network in which all nodes are within one hop of one another[24]. We used the implementation in `global_efficiency(G)` function from `NetworkX`[26].

**Near-criticality: Spectral radius.** We use the spectral radius as the representative near-criticality property. It is the largest eigenvalue magnitude of the adjacency matrix, $\rho(A) = \max_i |\lambda_i(A)|$, where $\lambda_i$ are the (possibly complex) eigenvalues of $A$. The spectral radius decides whether activity grows or dies out: above $1$ the dynamics amplify, below $1$ they decay. For ESNs it must therefore sit close to $1$ but below it, because higher values make the dynamics unstable and lower values cost expressivity. In brain networks the same quantity indexes how close the system is to a phase transition, and prior work discusses the assumption that brain networks sit at the edge of chaos, slightly below $\rho \approx 1$[11].

#### All 29 properties: definitions and landscapes

This section includes for each property level one table detailing all properties and one heatmap per property over the $25,000$ synthetic GNMs.

#### Goals: economy, capability, robustness

Table S2 lists the properties that read out the three goals - economy, capability, and robustness - together with each metric's definition, source, and formula. Figure S1 then maps every goal property across the $\eta$-$\gamma$ landscape of the $25{,}000$ synthetic networks.

Table S2: **Goal properties.** The goal-readout properties (economy, capability, robustness) evaluated for the 25,000 synthetic connectomes. The *Source* column gives the canonical reference for each metric's definition.

| Metric | Source | Why useful | Formula |
|---|---|---|---|
| **Economy** (be cheap) | | | |
| Wiring Cost | [6] | Physical cost of building the network: longer connections are more metabolically expensive. Brain networks are thought to minimise wiring cost under functional constraints. | $W = \sum_{ij} A_{ij} d_{ij}$ |
| Long-Range Fraction ($f_{\mathrm{LR}}$, at $39.56\%$) | [5] | Reliance on physically expensive long-range edges vs. short-range wiring, evaluated at the $39.56$th distance percentile (the threshold that best tracks absolute wiring cost). High values indicate high wiring cost. | $f_{\mathrm{LR}} = \frac{\lvert\{(i,j)\in E : d_{ij} > Q_q\}\rvert}{\lvert E\rvert}$, $Q_q$ = $q$-th quantile of all pairwise distances. |
| **Capability** (be able) | | | |
| Memory Capacity / IPC (linear, non-linear) | [10, 9] | Decomposes reservoir capacity by polynomial degree: linear (memory recall) vs. quadratic/cubic (nonlinear computation), separating the performance on delayed recall tasks from nonlinear computation performance. | $\mathrm{IPC}_{d,l} = (P z_{d,l})^2 / \lVert z_{d,l}\rVert^2$; $\mathrm{IPC}_{\mathrm{deg}_d} = \sum_l \mathrm{IPC}_{d,l}$. |

*continued on next page*

| Metric | Source | Why useful | Formula |
|---|---|---|---|
| Critical Dynamics (critical $T$, size, diversity) | [12] | Diversity of activity patterns the network spontaneously generates near criticality. More unique patterns indicate a richer repertoire of brain states; the count peaks near the critical threshold. | Cellular-automaton transitions; repertoire size $= |\mathcal{P}(T)|$ (unique patterns), $T_c = \arg\max_T |\mathcal{P}(T)|$. |
| Departure from Normality | [27] | Non-normality of $A$, which governs transient amplification: even a stable network can show large transients if highly non-normal. A capability readout of the spectral structure. | $d_F(A) = \|T_{\text{off}}\|_F / \|A\|_F$, $A = QTQ^*$ Schur, $T_{\text{off}} = T - \text{diag}(T)$. |
| **robustness** - be durable | | | |
| Robustness (targeted attack, and targeted/random ratio) | [13, 14] | How resilient the network is to node removal. Hub-dominated networks collapse quickly under targeted attack but resist random failure; the targeted-attack AUC and the targeted/random ratio index this classic trade-off. | $R = \frac{1}{N}\sum_{q=0}^{N-1} S_q$, $S_q = |LCC|/N$ after removing top-$q$ degree (targeted) or $q$ random nodes; $r = R_{targ}/R_{rand}$. |

How these properties are measured is set out in Section 2: the reservoir-computing setup and the information processing capacity (IPC) framework behind memory capacity, the cellular-automaton model behind the critical dynamics, and the node-removal experiment behind robustness. The figures there support each in turn. Figure 7 justifies the fraction of long-range connections $f_{\text{LR}}$ as a parcellation-invariant proxy for absolute wiring cost $W$, and the percentile threshold chosen for it. Figures 7, 8, and 8 compare memory capacity, critical dynamics, and attack tolerance across datasets.

#### Architecture: reach, centralization, segregation

Table S3 lists the properties that reads out the three architectural settings - reach, centralization, and segregation - together with each metric's definition, source, and formula. Figure S2 then maps every architecture property across the $\eta$-$\gamma$ landscape of the $25{,}000$ synthetic networks.

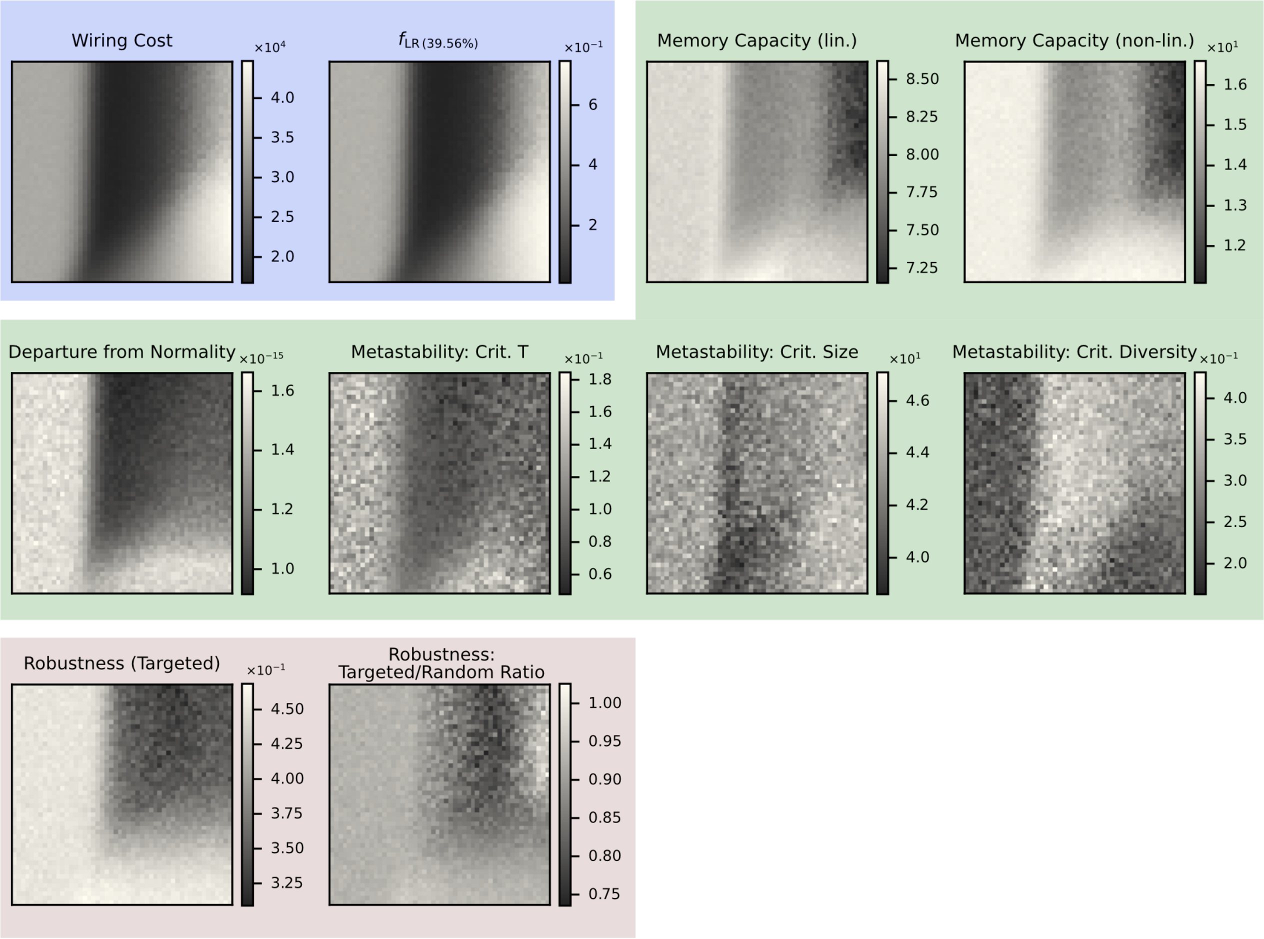


Figure S1: **Property landscapes of the goals.** Each panel maps one property across the $50 \times 50$ $\eta$-$\gamma$ grid (averaged over the 10 replicates per cell; $\eta$ increases along the horizontal axis, $\gamma$ along the vertical; darker colours indicate lower values). Panels cover every goal property that enters the property space: *economy* **- Wiring Cost** and the **Fraction of Long-Range Connections** ($f_{\text{LR}}$, $39.56$th percentile); *capability* **- Linear** and **Non-linear Memory Capacity** (IPC degree-1 and degree-2), critical dynamics **Critical** $T$, **Critical Repertoire Size**, **Critical Repertoire Diversity**, and **Departure from Normality**; *robustness* **- Robustness to Targeted Attack** (AUC) and the **Ratio of Targeted-to-Random-Attack Robustness**.

---

*continued on next page*

| Metric | Source | Why useful | Formula |
|---|---|---|---|

Table S3: **Architecture properties.** The architecture properties (reach, centralization, segregation) evaluated for the 25,000 synthetic connectomes.

| Metric | Source | Why useful | Formula |
|---|---|---|---|
| **reach** - edge length | | | |
| Ollivier-Ricci Curvature (min, skewness) | [15;18] | Local geometry per edge: positive curvature indicates parallel paths, negative curvature indicates a bottleneck or bridge. The minimum and skewness of the edge-curvature distribution capture reach (how far edges bridge) at mesoscale, missed by global measures. | $\kappa(x,y) = 1 - \frac{W_1(\mu_x^\alpha,\mu_y^\alpha)}{d(x,y)}$, $W_1$ = Wasserstein-1 distance between lazy-random-walk distributions. |
| **centralization** - hubs | | | |
| Rich-Club Coefficient ($k$ at max) | [28;29] | Whether high-degree hubs are disproportionately connected to each other ('rich club's), and a readout of centralization; reported as the degree $k$ at which the normalised coefficient peaks. | $\phi(k) = \frac{2E_{>k}}{N_{>k}(N_{>k}-1)}$; normalised $\phi_{\text{norm}}(k) = \phi(k)/\langle\phi_{\text{rand}}(k)\rangle$, $>1$ = rich-club. |
| Connector-hub fraction | [30] | Fraction of nodes that act as connector hubs ($P \approx 1$), bridging many modules; indexes how concentrated the integrative hub roles are. | $P_i = 1 - \sum_s (k_{is}/k_i)^2$; connector hubs have high $P_i$. Connector hubs are utilized as defined in [30]; the variation of utilizing the fraction of connector hubs is novel. |
| Average Controllability (std) | [19;20] | How easily a node can push the whole-brain system to nearby states. High-$\bar{\kappa}$ nodes are 'drivers' of state transitions; the standard deviation across nodes indexes how concentrated (centralized) control is. | $\bar{\kappa}_i = \text{tr}(B_i W_c B_i^\top)$, $W_c$ = controllability Gramian; $A$ normalised by $\lambda_{\text{max}}(A) + 1$. |

*continued on next page*

| Metric | Source | Why useful | Formula |
|---|---|---|---|
| **segregation** - modules | | | |
| Modularity | 21; 22 | Measures how cleanly the network divides into communities with dense internal and sparse external connections. High modularity indicates segregated, specialised subnetworks. | $Q = \frac{1}{2m}\sum_{ij}\left[A_{ij} - \frac{k_i k_j}{2m}\right]\delta(c_i, c_j)$; communities maximise $Q$ via Louvain. |
| Number Communities | 21; 22 | Number of modules in the modularity-maximising Louvain partition; a coarse count of how many segregated subnetworks the connectome resolves into. | $N_M = \lvert\{c_i\}\rvert$, the number of distinct community labels in the modularity-maximising partition. |
| Omega | 31 | Small-world index: $\omega = 0$ indicates small-world, $-1$ lattice, $+1$ random, summarising the lattice-to-random balance in one number. | $\omega = \frac{L_{rand}}{L} - \frac{C}{C_{latt}}$ |
| Directed Simplices (count, max size) | 32 | Counts higher-order feedforward motifs. For undirected graphs this reduces to counting maximal cliques; more simplices indicate more higher-order routing options. | Maximal ordered sets $(v_0, \ldots, v_n)$ with $A_{v_i v_j} = 1$ for all $i < j$. |
| Participation Coefficient (std) | 30 | How evenly a node's connections spread across communities. The spread (std) across nodes indexes how heterogeneous the connector- vs. provincial-hub roles are, a signature of modular segregation. | $P_i = 1 - \sum_s (k_{is}/k_i)^2$. |

#### Emergent behaviour: communication and near-criticality

Table S4 lists the properties of the two emergent behaviours - communication and near-criticality - together with each metric's definition, source, and formula. Figure S3 then maps every emergent-behaviour property across the $\eta$-$\gamma$ landscape of the $25{,}000$ synthetic networks.

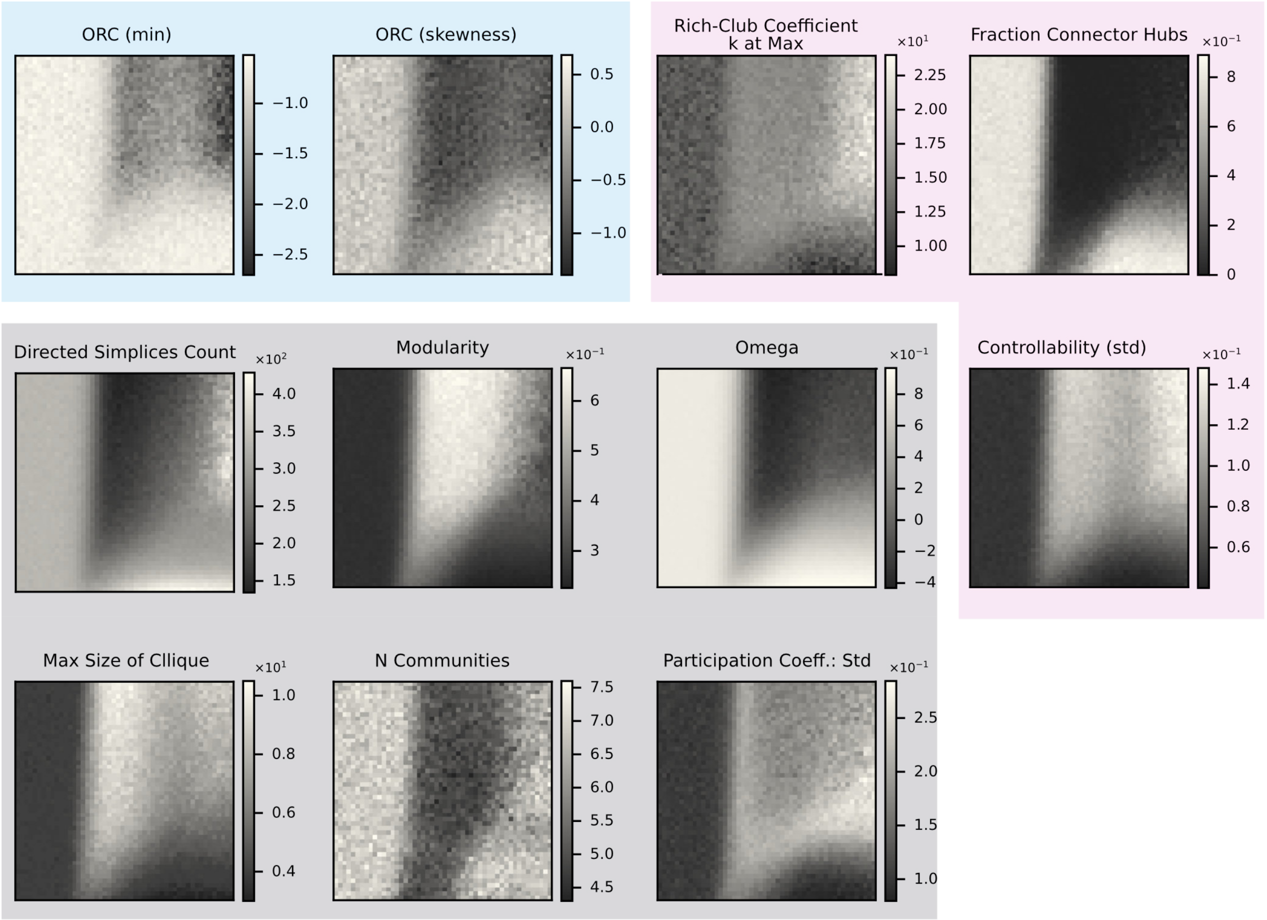


Figure S2: **Property landscapes of the architecture.** Each panel maps one property across the $50 \times 50$ $\eta$-$\gamma$ grid (averaged over the 10 replicates per cell; $\eta$ increases along the horizontal axis, $\gamma$ along the vertical; darker colours indicate lower values). Panels cover every architecture property that enters the property space: *reach* - **Ollivier-Ricci Curvature Minimum** and **Skewness**; *centralization* - **Rich-Club Coefficient** ($k$ at max), **Connector Hub Fraction**, and **Controllability Std** (NCT-STD); *segregation* - **Modularity**, **N Communities**, **Omega** ($\omega$), **Directed Simplices Count**, **Max Size of Clique**, and **Participation Coefficient std**.

Table S4: **Emergent-behaviour properties.** The communication and near-criticality properties evaluated for the 25,000 synthetic connectomes.

| Metric | Source | Why useful | Formula |
|---|---|---|---|
| **communication** - signalling | | | |
| Global Efficiency | 24 | Average inverse shortest-path length. How efficiently information flows between all node pairs in parallel; well-defined for disconnected graphs. | $E = \frac{1}{N(N-1)} \sum_{i \neq j} \frac{1}{d_{ij}}$ |

*continued on next page*

| **Metric** | **Source** | **Why useful** | **Formula** |
|---|---|---|---|
| Diffusion Efficiency | 33 | Efficiency of information diffusion via random walks (mean first-passage time), more biologically plausible than shortest paths for neural signalling. | $\mathrm{DE} = \frac{1}{N(N-1)} \sum_{i \neq j} H_{ij}^{-1}$, $H_{ij}$ = mean first-passage time. |
| Propagation Efficiency | 34 | How easily signals propagate via a communicability-style model: signals reach all nodes via many parallel walks without exponential attenuation. | $\mathrm{PE} = 1/\bar{\delta}$, $\delta_{ij} = -\log[(I-\alpha A)^{-1}(I-\alpha A)^{-\top}]_{ij}$, $\alpha = 0.8/\rho(A)$. |
| **near-criticality** - dynamics | | | |
| Spectral Radius | 11 | Largest eigenvalue magnitude of $A$. For reservoirs it must be $< 1$ for stability; for brain networks it indexes proximity to a dynamical phase transition. | $\rho(A) = \max_i \lvert\lambda_i(A)\rvert$ |
| Spectral Gap (Eigengap between largest eigenvalues) | - | Gap between the two largest eigenvalue magnitudes. A large gap means the leading mode dominates; relates to random-walk mixing time. In other fields known as "eigengap". | $\Delta = \lvert\lambda_1(A)\rvert - \lvert\lambda_2(A)\rvert$ |
| Synchronizability Eigenratio | 35 | Smaller $R$ = the network synchronises over a wider coupling range. Directly predicts whether coupled oscillators will synchronise. | $R = \lambda_N(L)/\lambda_2(L)$, $L = D - A$. |
| Fiedler Value / Algebraic Connectivity (and Laplacian spectral gap) | 36 | Bottleneck of diffusion and synchronisation: how robustly connected the network is. The $\lambda_2/\lambda_3$ gap (the Laplacian spectral gap) indicates a tendency toward exactly two communities. | $\lambda_2(L) = \min_{x \perp \mathbf{1}} \frac{x^\top L x}{x^\top x}$; normalised $\hat{\lambda}_2 = \lambda_2/\lambda_{\max}$. |

**Diffusion efficiency and propagation efficiency** Diffusion efficiency ($E_{\text{diff}}$) models information spread as an unbiased random walk and is defined as the inverse of the average commute time (expected number of steps for a random walker to travel from node $i$ to node $j$ and back)[37]. This requires no global topological knowledge and thus models decentralised signalling. Propagation efficiency ($E_{\text{prop}}$) is operationalised via communicability[34]: the $(i, j)$ entry of the matrix exponential $e^A$

(where $A$ is the adjacency matrix) counts all walks between $i$ and $j$, weighted by the inverse factorial of their length, giving a broadcasting-like measure that rewards networks with many short parallel pathways. Network-level $E_{\text{prop}}$ is computed as the mean off-diagonal entry of the communicability matrix, normalised to the fully connected graph. Both measures were implemented following[38] and cross-validated against the `netneurotools` library[39].

**Fiedler value (algebraic connectivity)** The Fiedler value $\lambda_2$ is the second-smallest eigenvalue of the combinatorial graph Laplacian $L = D - A$, where $D$ is the diagonal degree matrix and $A$ is the adjacency matrix[36]. It measures algebraic connectivity: $\lambda_2 = 0$ if and only if the graph is disconnected; larger values indicate that the network is more robustly connected and that random walks mix more rapidly. In the property space, $\lambda_2$ loads positively on PC1 alongside global efficiency, reflecting the co-alignment of integration and connectivity robustness. Computed via `numpy.linalg.eigh` applied to the Laplacian; disconnected networks return $\lambda_2 = 0$.

**Synchronizability eigenratio** The synchronizability eigenratio is defined here as $Q = \lambda_N / \lambda_2$, where $\lambda_N$ is the largest and $\lambda_2$ is the second-smallest eigenvalue of the graph Laplacian. This differs from the traditional definition of the synchronizability eigenratio (e.g.[35]). Networks synchronise more readily when $Q$ is small: a compact Laplacian spectrum makes it easier to find a coupling strength placing all modes in a stable synchronous regime. Computed from the same eigendecomposition as the Fiedler value.

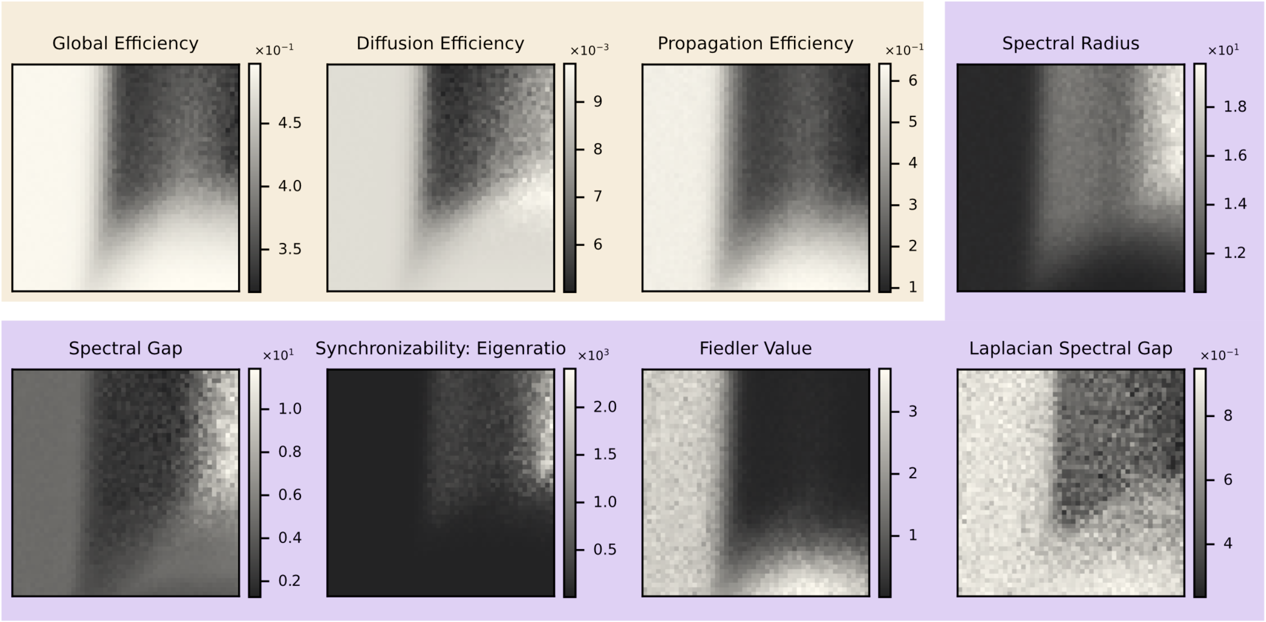


Figure S3: **Property landscapes of emergent behaviour.** Each panel maps one property across the $50 \times 50$ $\eta$-$\gamma$ grid (averaged over the 10 replicates per cell; $\eta$ increases along the horizontal axis, $\gamma$ along the vertical; darker colours indicate lower values). Panels cover every emergent-behaviour property that enters the property space: *communication* - **Global Efficiency** ($E_{\text{Global}}$, routing), **Diffusion Efficiency**, and **Propagation Efficiency** ($E_{\text{Prop}}$); *near-criticality* - **Spectral Radius**, **Spectral Gap**, **Synchronizability Eigenratio**, **Fiedler Value**, and **Laplacian Spectral Gap**.

#### Property Space: Additional Visualisations

Figure S4 extends the property-space analysis of the main text, colouring the PC1-PC2 morphospace by one structural property at a time.

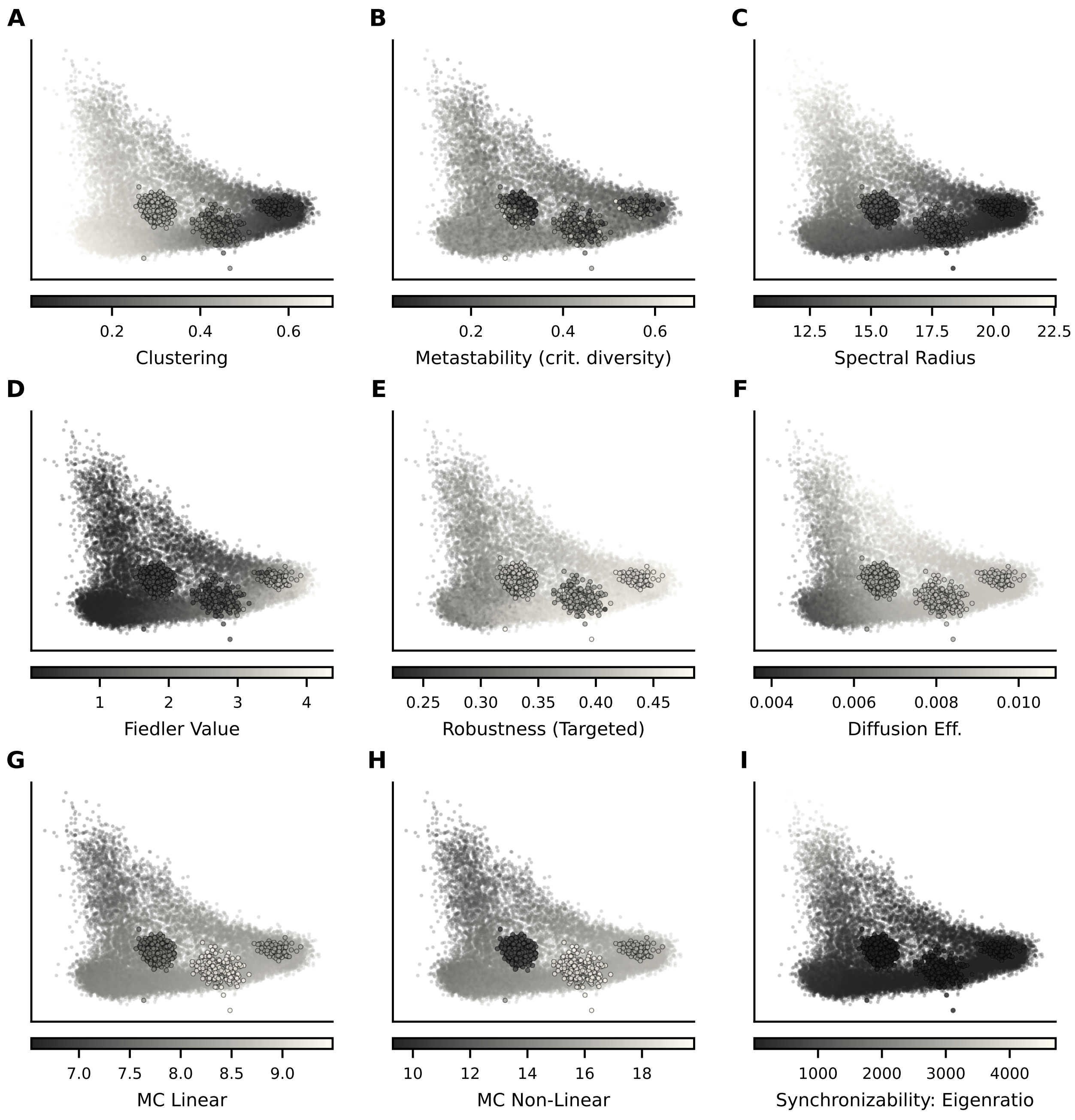


Figure S4: **The PC1-PC2 morphospace coloured by additional structural properties.** Each panel reproduces the PC1-PC2 morphospace with all GNM networks (small dots) and empirical, reference, and baseline networks (larger circles with black outline) coloured by the value of a single structural property.

#### Expansion of the Trade-Off Analyses

The ternary trade-off of the main text (Figure 3C, Section 2.2) normalises away the overall magnitude of its three axes and keeps only their relative balance. Figure S5 restores that magnitude as an orthogonal height axis, turning the triangle into the floor of a triangular prism, and plots the same networks as the main text.

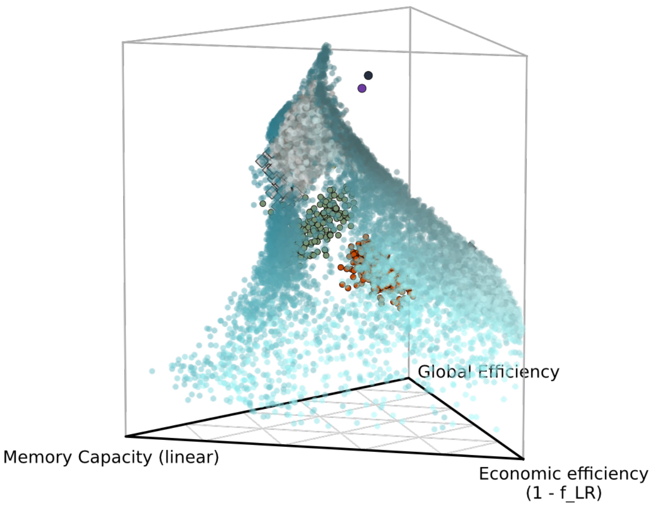


Figure S5: **Trade-off prism: relative balance (floor) and total rank (height) across economic efficiency, global efficiency, and memory capacity.** The triangular floor is the rank-normalised ternary plot of the main text (Figure 3C): for economic efficiency ($1 - f_{\mathrm{LR}}$), global efficiency, and linear memory capacity (the standalone forgetting-curve estimate at input scaling $s = 0.1$, not the $\mathrm{IPC}_1$ used elsewhere; see Methods), each network's value on an axis is replaced by its empirical-CDF position (rank) within the pooled distribution of all plotted networks, and the three ranks are divided by their sum, so floor position shows only the *relative* balance among the three objectives. The ternary projection discards how high the three ranks are *together*; the prism restores this as an orthogonal height axis $z$ equal to the summed rank (range $[0, 3]$), so a point's height is its *total* standing across all three objectives at once. No network reaches the memory-capacity vertex of the floor, and empirical connectomes (black outline) sit at intermediate heights, below the low-$\gamma$ GNM networks that attain the greatest total rank. The same 25,000 GNM networks (teal), empirical connectomes (orange: HCP-D; olive: MaMI), structural baselines (Erdős-Rényi; minimum-wiring network), and communication-optimised reference networks (coloured markers) are shown.

#### GNM-Fitted Property Space: PCA Loadings

The main text reads the GNM-fitted property space through its first two components (Section 2.4). The full loadings across all 29 properties, for the first three principal components, are given below.

#### Sensitivity of the Developmental Inflection Age to Smoothing Parameter

To assess the robustness of the identified developmental inflection, the curvature analysis (Equation 2) was repeated across smoothing parameters $\lambda \in \{1, 2, \ldots, 20\}$ and the generalised cross-validation (GCV) default (`lambda = None` in `scipy.interpolate.make_smoothing_spline`). Figure S7 shows the age of maximum spline curvature as a function of $\lambda$. The inflection age is stable at 16.0 years for all $\lambda \geq 5$ (16 of the 20 integer values tested). For $\lambda \in \{2, 3, 4\}$ and the GCV default ($\lambda \approx 2$), an earlier inflection near 9.5 years is identified, corresponding to a minor local bend in the less-smoothed trajectory that is suppressed once sufficient smoothing is applied. We adopted $\lambda = 10$ as the primary value: it lies well within the stable plateau while avoiding excessive over-smoothing.

#### Phylogenetic Order in the GNM Morphospace

The PC1-PC2 property space cleanly separates all five mammalian orders (PERMANOVA pseudo-$F \approx 16$ on PC1; Section 2.4). Embedding the same connectomes in the GNM $(\eta, \gamma)$ morphospace via DeltaCon best-fit parameters substantially reduces that separation. Pairwise group comparisons between most taxonomic orders are non-significant there (Table S5 and Figure S8). The

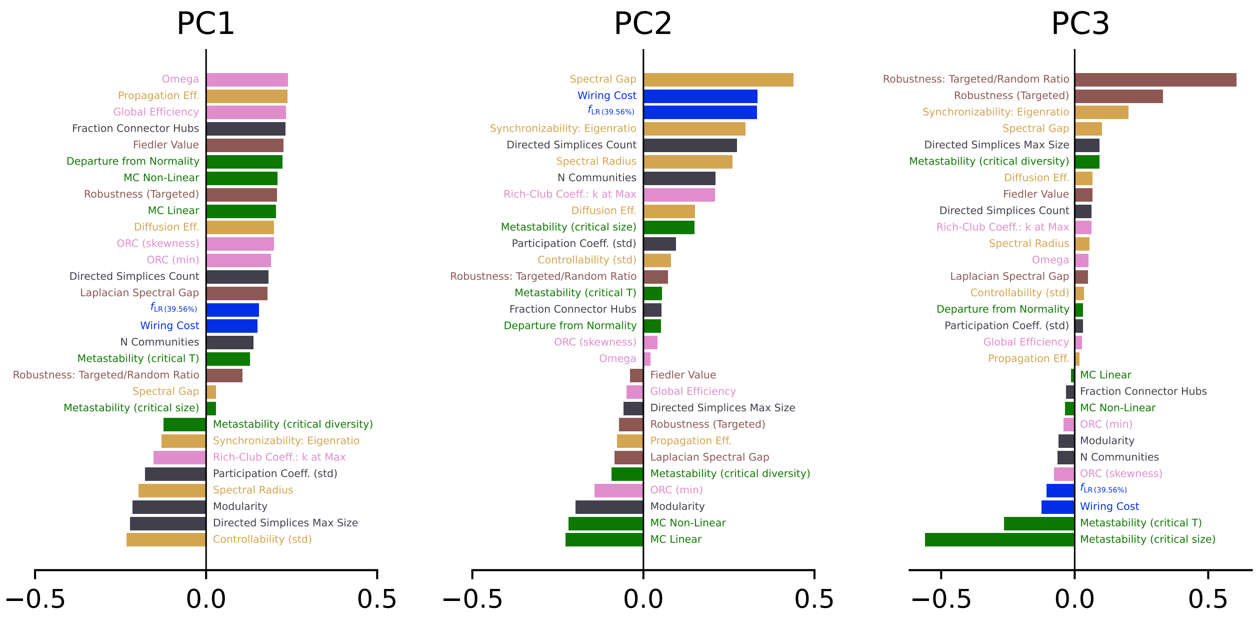


Figure S6: **PCA loadings for the GNM-fitted property space (full detail).** Loadings of the first three principal components of the PCA fit only on GNM networks, showing the contribution of all 29 network properties.

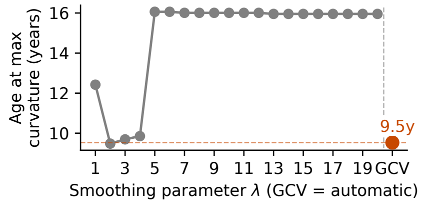


Figure S7: **Sensitivity of the developmental inflection age to the spline smoothing parameter.** Age of maximum trajectory curvature as a function of $\lambda$. The GCV-selected value (orange dot, $\lambda \approx 2$) identifies an inflection near 9.5 years; values $\lambda \geq 5$ consistently yield an inflection at $\approx 16.0$ years. The primary analysis uses $\lambda = 10$, as this is within the stable plateau.

reason is that the two-dimensional $(\eta, \gamma)$ space parameterises only the wiring-cost against homophily trade-off. It compresses the full 29-dimensional property variation onto a single axis, and loses the orthogonal variation that distinguishes mammalian orders in the property space. The GNM morphospace is therefore informative about *where* connectomes fall on the economy-homophily spectrum, but not about their phylogenetic identity.

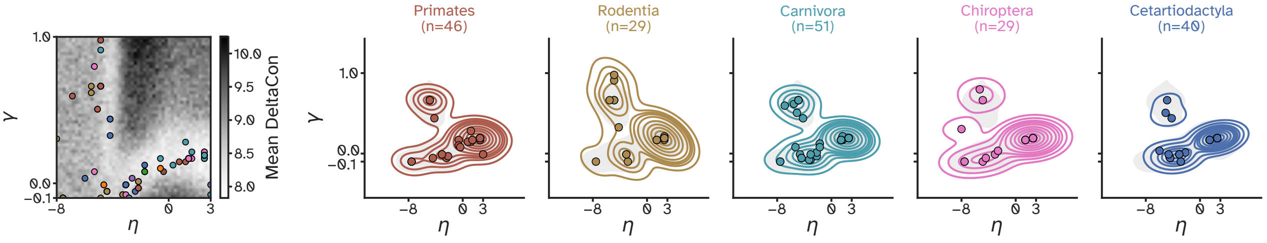


Figure S8: **Mammalian connectomes coloured by taxonomic order in the GNM $(\eta, \gamma)$ morphospace.** Each point shows the best-fitting GNM parameter combination for one MaMI connectome (identified via DeltaCon), coloured by taxonomic order; orders with $n > 20$ are highlighted. PERMANOVA results on these coordinates are reported in Table S5.

| **Order A** | **Order B** | $F$ | $p$ | Cohen's $d$ |
|---|---|---|---|---|
| Primates | Rodentia | 0.01 | 0.955 ns | 0.01 |
| Primates | Carnivora | 0.49 | 0.478 ns | 0.14 |
| Primates | Chiroptera | 0.00 | 0.964 ns | 0.02 |
| Primates | Cetartiodactyla | 0.02 | 0.887 ns | 0.03 |
| Rodentia | Carnivora | 0.35 | 0.547 ns | 0.14 |
| Rodentia | Chiroptera | 0.01 | 0.953 ns | 0.01 |
| Rodentia | Cetartiodactyla | 0.04 | 0.841 ns | 0.03 |
| Carnivora | Chiroptera | 0.27 | 0.609 ns | 0.12 |
| Carnivora | Cetartiodactyla | 0.68 | 0.420 ns | 0.18 |
| Chiroptera | Cetartiodactyla | 0.03 | 0.869 ns | 0.04 |

Table S5: **Phylogenetic order separation is lost in the GNM morphospace.** Pairwise PERMANOVA (999 permutations) on the best-fitting $(\eta, \gamma)$ GNM coordinates assigned to each MaMI connectome via DeltaCon. All pairwise group comparisons are non-significant ($p \geq 0.05$), indicating that the two-dimensional wiring-rule parameterisation does not preserve the phylogenetic structure visible in the property space. $F$: pseudo-$F$ statistic. $p$: permutation $p$-value. Cohen's $d$ computed on the $\eta$ coordinate. Significance: *** $p < 0.001$, ** $p < 0.01$, * $p < 0.05$, ns $p \geq 0.05$.

[4] Kayson Fakhar, Danyal Akarca, Andrea I. Luppi, Stuart Oldham, Fatemeh Hadaeghi, Petra E. Vértes, Ed Bullmore, Claus Hilgetag, and Duncan Astle. Human cortical networks trade communication efficiency for computational reliability, December 2025.

[5] Stuart Oldham, Alex Fornito, and Gareth Ball. Coming up short: Generative network models fail to accurately capture long-range connectivity, November 2024.

[6] Ed Bullmore and Olaf Sporns. The economy of brain network organization. *Nature Reviews. Neuroscience*, 13(5):336–349, April 2012.

[7] H. Jaeger. Short term memory in echo state networks. 2001.

[16] Chien-Chun Ni, Yu-Yao Lin, Jie Gao, Xianfeng David Gu, and Emil Saucan. Ricci Curvature of the Internet Topology, January 2015.

[17] Hamza Farooq, Yongxin Chen, Tryphon T. Georgiou, Allen Tannenbaum, and Christophe Lenglet. Network curvature as a hallmark of brain structural connectivity. *Nature Communications*, 10(1):4937, October 2019.

[18] Chien-Chun Ni, Yu-Yao Lin, Feng Luo, and Jie Gao. Community Detection on Networks with Ricci Flow. *Scientific Reports*, 9(1):9984, July 2019.

[19] Shi Gu, Fabio Pasqualetti, Matthew Cieslak, Qawi K. Telesford, Alfred B. Yu, Ari E. Kahn, John D. Medaglia, Jean M. Vettel, Michael B. Miller, Scott T. Grafton, and Danielle S. Bassett. Controllability of structural brain networks. *Nature Communications*, 6(1):8414, October 2015.

[20] Linden Parkes, Jason Z. Kim, Jennifer Stiso, Julia K. Brynildsen, Matthew Cieslak, Sydney Covitz, Raquel E. Gur, Ruben C. Gur, Fabio Pasqualetti, Russell T. Shinohara, Dale Zhou, Theodore D. Satterthwaite, and Dani S. Bassett. Using network control theory to study the dynamics of the structural connectome, August 2023.

[39] Zhen-Qi Liu, Vincent Bazinet, Justine Y. Hansen, Filip Milisav, Andrea I. Luppi, Eric G. Ceballos, Asa Farahani, Laura E. Suarez, Golia Shafiei, Ross D. Markello, and Bratislav Misic. Netneurotools: A trainee-oriented approach to network neuroscience, September 2025.
